# Commensal bacteria act as a broad genetic buffer in *Drosophila* during chronic under-nutrition

**DOI:** 10.1101/334342

**Authors:** Dali Ma, Maroun Bou-Sleiman, Pauline Joncour, Claire-Emmanuelle Indelicato, Michael Frochaux, Virginie Braman, Maria Litovchenko, Gilles Storelli, Bart Deplancke, François Leulier

**Author notes:** Present address: Laboratory of Integrative Systems Physiology, Interschool Institute of Bioengineering, School of Life Sciences, Ecole Polytechnique Federale de Lausanne (EPFL) CH-1015, Lausanne, Switzerland. Present address: Department of Human Genetics, University of Utah School of Medicine, Salt Lake City, UT 84112, USA.

## Abstract

Eukaryotic genomes encode several well-studied buffering mechanisms that robustly maintain invariant phenotypic outcome despite fluctuating environmental conditions. Here we show that the gut microbiota, represented by a single Drosophila facultative symbiont, *Lactobacillus plantarum* (*Lp*^*WJL*^), acts also as a broad genetic buffer that masks the contribution of the cryptic genetic variations in the host under nutritional stress. During chronic under-nutrition, *Lp*^*WJL*^ consistently reduces variation in different host phenotypic traits and ensures robust organ patterning; *Lp*^*WJL*^ also decreases genotype-dependent expression variation, particularly for development-associated genes. We further demonstrate that *Lp*^*WJL*^ buffers via reactive oxygen species (ROS) signaling whose inhibition severely impairs microbiota-mediated phenotypic robustness. We thus identified an unexpected contribution of facultative symbionts to *Drosophila* fitness by assuring developmental robustness and phenotypic homogeneity in times of nutritional stress.

## Results and Discussions

### Mono-association with Lp^WJL^ reduces growth/size variation of Drosophila larvae during chronic under-nutrition in the DGRP lines

Despite environmental stress, organisms possess intrinsic genetic buffering mechanisms to maintain phenotypic constancy by repressing the expression of cryptic genetic variants, thus preserving genetic diversity. Compromising these buffering mechanisms unlocks new substrates for natural selection[1-3]. However, natural selection can also operate on the hologenome, as symbiosis is recognized as a major driving force of evolution [4, 5]. Facultative nutritional mutualism forged by the host and its resident gut microbiota permits the holobiont to adapt to changing nutritional environments during the host’s life time[6]. Consequently, the evolutionary implications of such association deserve more scrutiny. Horizontally acquired gut bacteria in *Drosophila* are a perfect example of nutritional mutualists[7]. Previously, we showed that a single commensal strain, *Lp*^*WJL*^ can significantly accelerate growth in ex-germ free (GF) larvae during chronic undernutrition[8, 9]. To study the host’s genetic contribution to *Lp*^*WJL*^-mediated growth in the same context, we first measured the body lengths of both the GF and *Lp*^*WJL*^ mono-associated larvae from 53 DGRP lines 7 days after post-embryonic development (Fig.1a-c; Table S1), and conducted genome-wide association studies (GWAS) based on the ranking of growth gain by comparing GF and *Lp*^*WJL*^-associated animals (Fig.1a; TableS1, column “ratio”). The GWAS yielded nine candidate variants (Table S2, Fig.S1a and S1b), and through RNA interference (RNAi), we assessed the contribution of each variant-associated gene to host growth with or without *Lp*^*WJL*^. Surprisingly, we failed to capture any obvious “loss or gain of function” of the growth benefit conferred by *Lp*^*WJL*^. Instead, we observed that the individual RNAi-mediated knock-down of gene expression led to large phenotypic variation in GF larvae, but such variation was reduced in *Lp*^*WJL*^, resulting in growth gain in all tested genetic backgrounds (Fig.S1c and S1d). In parallel, we computed the respective heritability estimates (H) for the GF and *Lp*^*WJL*^-associated DGRP populations as 37% vs. 10% (Fig.1b and 1c). The coefficient of variation (CV) of the pooled GF population was also greater, despite their overall smaller standard deviation and average size (Fig.1d). These three observations strongly indicate that genetic variants induce more pronounced size variation in GF animals, and the gut bacteria unexpectedly restrict growth variation despite host genetic differences. Next, we plotted the individual average GF larval length value against that of its *Lp*^*WJL*^-associated siblings from both the DGRP and the RNAi studies and derived the linear regression coefficients. If genetic background predominantly impacts growth, then this coefficient should be close to 1, yet we found that both are close to zero (0.145 and 0.06 respectively; Fig.1e and Fig.S1e). We thus conclude that *Lp*^*WJL*^ presence effectively masks the contribution of genetic variation in the DGRP lines and steers the animals toward attaining a similar size independent of genotype.

**Figure 1.**
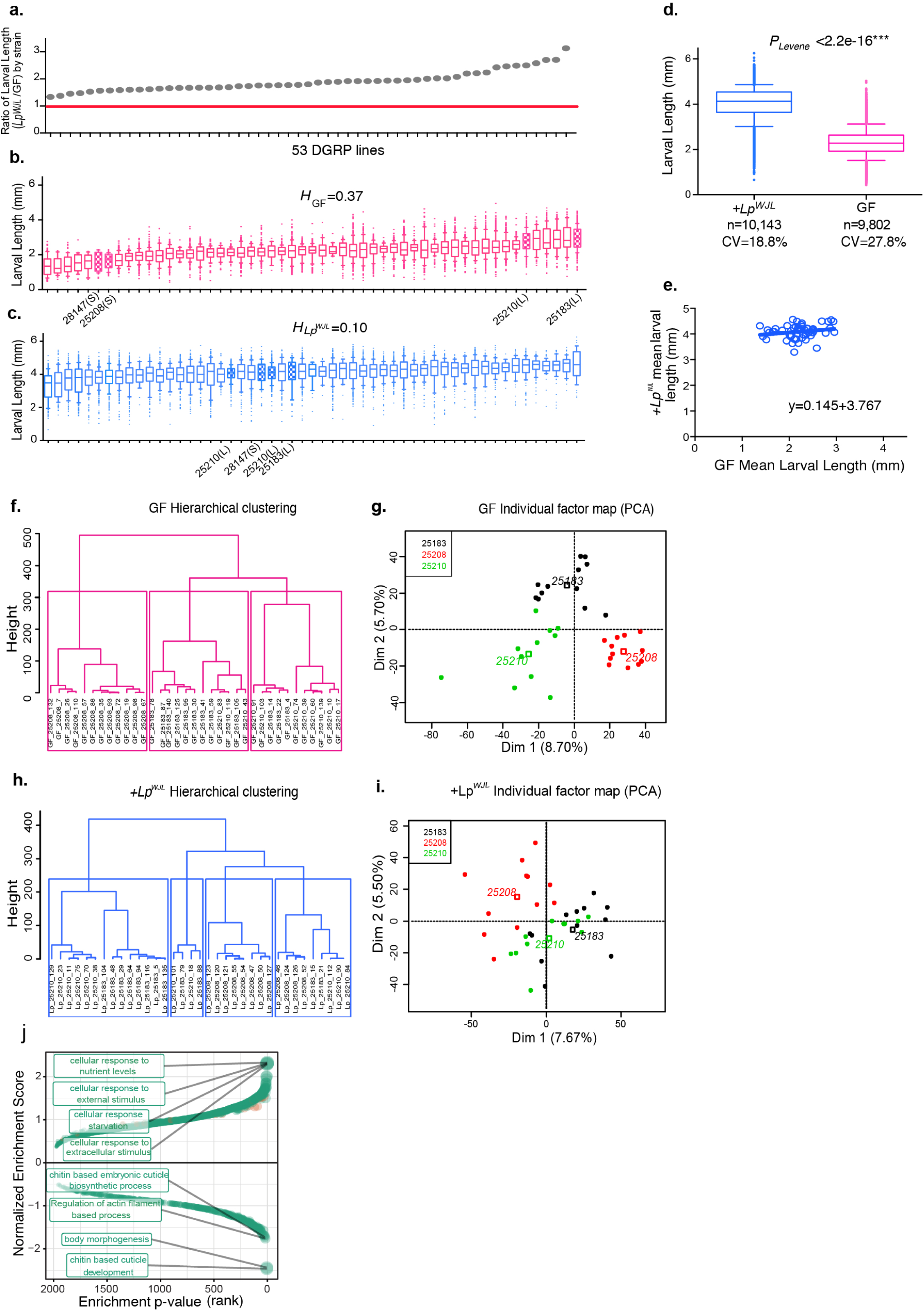
Mono-association with *Lp*^*WJL*^ buffers phenotypic and transcriptomic variation during growth and development in the DGRP lines. **a).**The ranking of larval growth gain of 53 DGRP lines was used for GWAS to uncover host variants associated with growth benefits conferred by *Lp*^*WJL*^. Each grey dot represents the quotient of average mono-associated larval length (Figure 1c) divided by the average length of GF larval length (Figure 1b) from each DGRP line on Day 7 AEL (<u>a</u>fter <u>e</u>gg lay). The red line marks the ratio of “1”, indicating that all tested DGRP lines benefited from *Lp*^*WJL*^ presence. **b).**and **c)**. the average larval length on Day 7 AEL for each of the 53 DGRP lines (Mean and 10-90 percentile. Unless specified, all box plots in this manuscript present the same parameters). Each line in the box represents the average length from pooled biological replicates containing all viable larvae from all experimental repeats. From each strain, there are between 10-40 viable larvae in each replicate, 3 biological replicates for each experiment, and 2 to 3 repeats of the experiments. **b):** germ-free (GF, pink), **c):** mono-associated (+*Lp*^*WJL*^, blue). Note the heritability estimate (H) in the GF population is higher than in the mono-associated population (37% vs. 10%). The filled boxes denote the “small (S)” and “large (L)” DGRP lines that were selected for setting up the F_2_ crosses (see Figure S3a for crossing schemes). **d).**Box and whiskers plots showing average larval length derived from pooled GF (pink) or *Lp*^*WJL*^- (blue) mono-associated DGRP lines. The coefficient of variation in the GF population (27.82 %) is greater than that of the mono-association population (18.74%). Error bars indicate 10 to 90^th^ percentile. Levene’s test is used to evaluate homocedasticity and Mann-Whitney test for difference in the mean (*P*<0.0001****). **e).**Scatter plot to illustrate that *Lp*^*WJL*^ buffers size variation in ex-GF larvae in the DGRP population. Each data point represents the intercept of the GF length and its corresponding mono-associated length at Day 7 for each DGRP line. If genetic variation was the only factor influencing growth in both GF and monoassociated flies, the slope of the scatter plot should theoretically be 1 (Null hypothesis: slope=1. *P<0.0001*: the null hypothesis is therefore rejected. A linear standard curve with an unconstrained slope was used to fit the data). **f) g)., h)**. and **i)**. Hierachical clustering (**f** and **h**) and PCA analyses (**g** and **i**) based on individual larvae transcriptome analyses show that the samples cluster more based on genotypes when germ-free (**f** and **g**, **f**: *P*_genotype_=1.048e-08, **g**: R^2^ _Dim1_=0.73, *P*_genotype_=7.81e-10, R^2^ _Dim2_=0.72, *P*_genotype_=1.12e-9,) than mono-associated (**h** and **i**, **h**: *P*_genotype_ =0.000263, **i:** R^2^ _Dim1_=0.42, *P*_genotype_ =0.00017**, R^2^ _Dim2_=0.31, *P*_genotype_=0.00269). A PCA followed by hierarchical clustering on principle components (HCPC) was performed with the R package FactoMineR on the voom corrected read counts. Correlations between the genotype variable and PCA dimensions or HCPC clusters were assessed by χ^2^ tests. The dots represent the different samples according to genotype, and the empty squares are the calculated centers for each genotype. **j)**. Gene set enrichment analysis based on the change in standard deviation of gene expression. Positive enrichment indicates gene sets that are enriched in the genes whose expression level variation increases in response to *Lp*^*WJL*^ mono-association. Negative gene sets are those that are enriched in the genes whose expression level variation decreases in response to *Lp*^*WJL*^ mono-association. The top 4 positively and negatively enriched sets are labeled. The genes whose expression levels are reduced by *Lp*^*WJL*^ mono-association predominantly act in chitin biosynthesis and morphogenesis (See also FigS2.

### Mono-association with Lp^WJL^ decreases expression variation of developmentally-related genes during growth

Since *Lp*^*WJL*^ reduces growth variation in the host phenotypically, and phenotypic variation is often the manifestation of transcriptomic variation due to genetic differences[10], we explored if *Lp*^*WJL*^ also decreases gene expression variation during larval development. We conducted BRB-seq[11] on 36 mono-associated and 36 GF individual larvae from 3 DGRP lines and specifically compared transcriptional variation in individual *Lp*^*WJL*^ mono-associated larvae to that of age-matched GF samples (Fig.S2a). First, we observed that the transcriptomes tend to cluster by genotype and *Lp*^*WJL*^ status after batch effect correction (Fig.S2b and S2c, Table S3). Second, the overall transcriptomic changes and the GO-terms associated with *Lp*^*WJL*^ presence corroborate our previous findings, as similar sets of digestive enzymes and immune response genes were up-regulated (Fig.S2d and S2e)[12]. Interestingly, genotype was a stronger clustering driver for GF samples than *Lp*^*WJL*^ mono-associated ones when samples were separated based on bacterial presence (Fig 1f vs. 1h, and 1g vs. 1i). These observations suggest that *Lp*^*WJL*^ can mask host genetic differences also at the transcriptomic level. Next, we compared the standard deviation (SD) of each expressed gene in both conditions, and found that even though mono-association can either elevate or reduce expression variation in different gene sets (Fig.S2f and S2g), there is a tendency toward an overall increase in expression variation in GF transcriptomes (Fig.S2f, red line). This trend was also more apparent in genes that were non-differentially expressed between the GF and mono-associated conditions (Fig.S1h, middle panel, grey lines). Finally, we found that genes whose expression variation most-decreased by *Lp*^*WJL*^ are enriched in developmental processes such as “body morphogenesis” and “cuticle development” (Fig.1j). These data reveal that *Lp*^*WJL*^ mono-association dampens genotype-dependent expression variation, especially of genes linked to developmental processes, which in turn may account for the ability of *Lp*^*WJL*^ to reduce larval size variation.

### Lp^WJL^ broadly buffers variation in different physical fitness traits in genetically diverse populations

Based on these findings, we propose that *Lp*^*WJL*^ effectively reduces both phenotypic and transcriptional fluctuations during chronic under-nutrition. *Lp*^*WJL*^ thus confers a biological function that resembles various canonical buffering mechanisms that robustly maintain phenotypic homogeneity by masking the effects of cryptic genetic variation[2, 13, 14], despite the presence of a persistent nutritional stress signal. Since our studies insofar were conducted only in homozygous inbred DGRP lines, we sought to test if the observed buffering also operates in a population of heterozygous and genetically diverse individuals. Therefore, based on their GF growth profile, we selected two DGRP strains from each end of the phenotypic extremes (Fig.1b and 1c, patterned pink and blue bars), established seven different inter-strains crosses, and compared the growth variation in the GF and mono-associated F_2_ progenies (Fig.S3a, Methods). In these experiments, we supplemented the GF larvae with 33% more yeast (8g.L^-1^ vs 6g.L^-1^) to address two possible caveats: first, we wished to exclude that *Lp*^*WJL*^ might simply act as an additional food source, even though our previous findings indicate that this is not the case[7]. However, if increasing the dietary yeast content reduces the variability in GF growth to the same extent as the gut bacteria, then the buffering effect may be generally attributed to mere superior food quality. Second, greater yeast content accelerates GF growth; consequently, the size and stage differences between the GF and mono-associated larvae are minimized, thus allowing us to compare variation in size-matched GF and mono-associated larvae *en masse*, while excluding the bias that bigger and older mono-associated larvae might vary less as they tend to be more mature.

Our hypothesis predicts that the GF F_2_ population should show higher variance in body length than their *Lp*^*WJL*^ mono-associated siblings. Indeed, in the F_2_ larvae, the CV and SD values tend to separate into two distinct groups, as driven by *Lp*^*WJL*^ presence (Fig.2a, Fig.S3b). Overall, the F_2_ *Lp*^*WJL*^ mono-associated larvae were slightly longer, but their GF siblings varied more in length, regardless of yeast content or larval age (Fig.S3c). In the size-matched pools (Fig.2a, purple bracket), GF size still fluctuated more than that of the *Lp*^*WJL*^ mono-associated siblings (Fig.2b), despite the fact that they were fed with a richer diet. Based on these observations, we first confirm that augmenting yeast content fails to recapitulate the same buffering effect mediated by living commensals. More importantly, we conclude that phenotypic buffering by the gut microbe *Lp*^*WJL*^ indeed operates in a genetically diverse host population facing a nutritional challenge, hence qualifying the gut microbiota as a previously unappreciated buffering agent of cryptic genetic variation.

**Figure 2.**
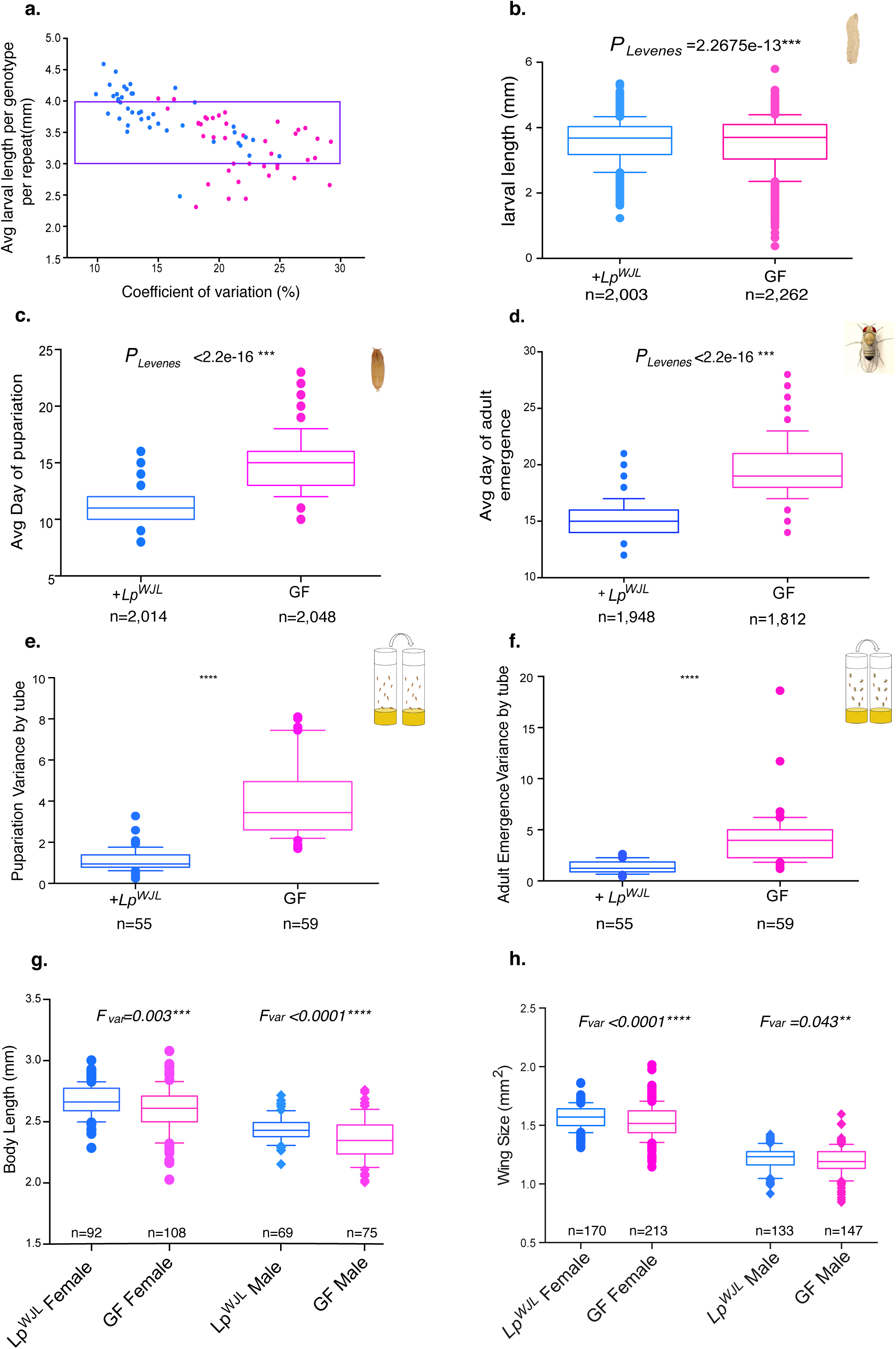
In the genetically diverse DGRP F_2_ population, *Lp*^*WJL*^ reduces variation in different physical fitness traits. **a).**A scatter plot showing how coefficient of variation (CV) changes as a function of larval length, and how such change differs in the DGRP F_2_ GF (pink) and *Lp*^*WJL*^ mono-associated (blue) populations (see Figure S3a and Methods for detailed schemes). Each data point represents the intercept of a CV value and its corresponding average larval length in a particular cross. Each CV, SD and average value was derived from larvae measurements gathered from at least 3 biological replicates from either GF or *Lp*^*WJL*^ mono-associated conditions. Each replicate contains 10-40 larvae. Based on multivariate anova analysis, the factors affecting variants in this plot are: larval age* (P=0.053), bacteria presence***(P=3.02e-06), and larval length (P=8.27e-15***). The purple bracket indicates the arbitrarily selected experiments where the average larval length for each cross falls between 3mm and 4mm for size-matching purpose. **b).**The average larval length of the F_2_ progeny pooled from experiments demarcated by the purple bracket in Figure **2a**. While the average size is perfectly matched (GF Avg Length=3.522mm, *Lp*^*WJL*^ Avg Length= 3.582mm, P=0.857^ns^, Mann-Whitney test), the GF population exhibits greater variation than the *Lp*^*WJL*^ mono-associated population (Var_GF_=0.642, CV_GF_=22.8%, Var_Lp_=0.427, CV_Lp_=18.3%) **c).**Variance and mean comparisons for the average day of pupariation for individual larva in the F_2_ GF and mono-associated populations. (Difference in mean P<0.0001***, Mann-Whitney test, Var _GF_= 2.42, Var_Lp_=1.22). **d).**Variance comparison for average day of adult emergence in the F_2_ GF and mono-associated populations (Difference in mean P<0.0001***, Var_Lp_=1.84, Var_GF_=5.27). **e).**Box plots comparing the variances of pupariation derived from each tube containing approximately 40 larvae. The average variance per tube for the GF population=3.99; the average variance per tube for the *Lp*^*WJL*^ associated population =1.12. Var_Lp_=0.54, Var_GF_=1.76. Note that these values are the “variance of variances”. **f).**Box plots comparing the variances for adult emergence from each tube containing approximately 40 larvae (Difference in mean P<0.0001***). The average variance per tube for the GF population=4.06; the average variance per tube for the *Lp*^*WJL*^ associated population =1.34. For ‘variance of the variances”, Var_Lp_=1.33, Var_GF_=4.2. **g)**. and **h.**) In both male (lozenge) and female (circle) adults, the variances in body size (**g**. the difference in mean body length: for females, P=0.0009***, for males, P=0.0015**), and wing size (**h**., the difference in mean wing area for females, P=0.0010, *** for males, P=0.124, ns) are greater in the GF population than in the mono-associated population. The adult data sets presented in Fig.**2g** and **2h** and in Fig.**S3g** and **S3h** take on normal distribution by D’Agostino & Pearson omnibus normality test, F variances are therefore calculated and compared.

During chronic under-nutrition, *Lp*^*WJL*^ sustains growth rate as effectively as an entire gut microbiota[8]. We thus wondered if a natural and more complex gut microbiota can also buffer growth variation like *Lp*^*WJL*^. To address this question, we rendered a population of wild flies collected in a nearby garden germ-free, and re-associated them with their own fecal microbial community[15]. In three out of four experimental repeats, growth variation is significantly reduced in the larval population fed on food inoculated with fecal microbiota (Fig.S3d and data not shown), and the cumulative CV and variances derived from each food cap were significantly higher in the GF population (Fig.S3e and S3f). This strongly suggests that the gut-associated microbial community of wild flies indeed decreases growth variation of a natural *Drosophila* population. However, since the wild-derived microbiota did not consistently buffer larval growth, probably due to the difficulty to precisely control the quantity and composition of the inoculated fecal microbiota, we returned to the mono-association model for subsequent studies.

If the observed growth variation in GF larvae indeed reflects the “unleashing” of the host’s genetic potential due to the loss of a buffering mechanism provided by gut microbes[2], then we hypothesized that other physical fitness traits in a fertile surviving GF population should in principle also exhibit greater phenotypic variation. We therefore examined the variances in pupariation timing and adult emergence in the F_2_ progeny of the inter-DGRP strain crosses (Fig.S3a). First, individual GF larvae pupariated and eclosed later, but the variances in the pooled data were greater than that of mono-associated counterparts (Fig.2c and 2d); from each vial containing an equal number of larvae, the variances of pupariation and eclosion were also greater in the GF samples (Fig.2e and 2f). Therefore, both inter-individual and among-population variances in developmental timing and adult emergence are reduced. Lastly, GF adults were slightly shorter (Fig.2g); the sizes of representative organs, expressed as area of the eye and the wing, were also smaller, yet the variances in these traits were greater (Fig.2h, Fig.S3g). Furthermore, the wing/body-length allometric slopes remained unaltered, but the individual GF values were more dispersed along the slope (Fig.S3i,j); when taken as a ratio (wing length/body-length), the variance was greater in the GF flies (Fig.S3h). These observations indicate that gut microbes, represented by *Lp*^*WJL*^, indeed act as a broad buffer that confers phenotypic homogeneity in various physical fitness traits in a genetically diverse host population.

### Lp^WJL^ conveys robustness in organ-patterning under nutritional stress

We have thus far shown that *Lp*^*WJL*^ association confers transcriptomic stability and phenotypic constancy to the developing host facing nutritional stress, in a fashion that is highly reminiscent of the host’s own genetic buffering mechanism. For example, reducing Hsp90 activity has been shown to increase organ size variation in both plants and animals[16-18]. Moreover, compromising Hsp90 can lead to morphological aberrations that are otherwise “hidden”[17]. Similarly, we also found that a significant fraction of the GF F_2_ flies bore aberrant wing patterns such as missing margins, incomplete vein formations and ectopic vein tissue (Fig.3a). The incidence of wing anomalies differed according to the genotype, and females were more affected than males (Fig.3b). In contrast, the most visible “defect” in their *Lp*^*WJL*^ associated siblings, if any, were rare and hardly discernable (Fig.3a, Fig.S4a). Furthermore, gross patterning anomalies were absent in the viable adults from the GF parental homozygous strains or in F_2_ adults reared on a standard diet (data not shown), supporting that gut microbiota likely acts as a developmental canalization mechanism by suppressing the contribution of cryptic genetic variation in the presence of nutritional stress. Organ patterning is a robust process; changes in nutrition, humidity, temperature and crowding can alter the final adult body and wing size; yet wing patterning is virtually invariant and reproducible[19]. Surprisingly, we found that in GF flies, constant nutritional stress can in fact unveil the effects of preexisting “silent” mutations that manifest themselves as visible wing patterning anomalies. Furthermore, as the patterning defects only appear in nutritionally challenged F_2_ flies devoid of their microbiota, we conclude that these defects reflect a breach of the canalization process during developmental patterning when the hidden effects of genetic variants are unlocked[20], and the gut microbiota buffers the effects of these otherwise seemingly “neutral” variants to confer robustness to the canalized process of organ patterning.

**Figure 3.**
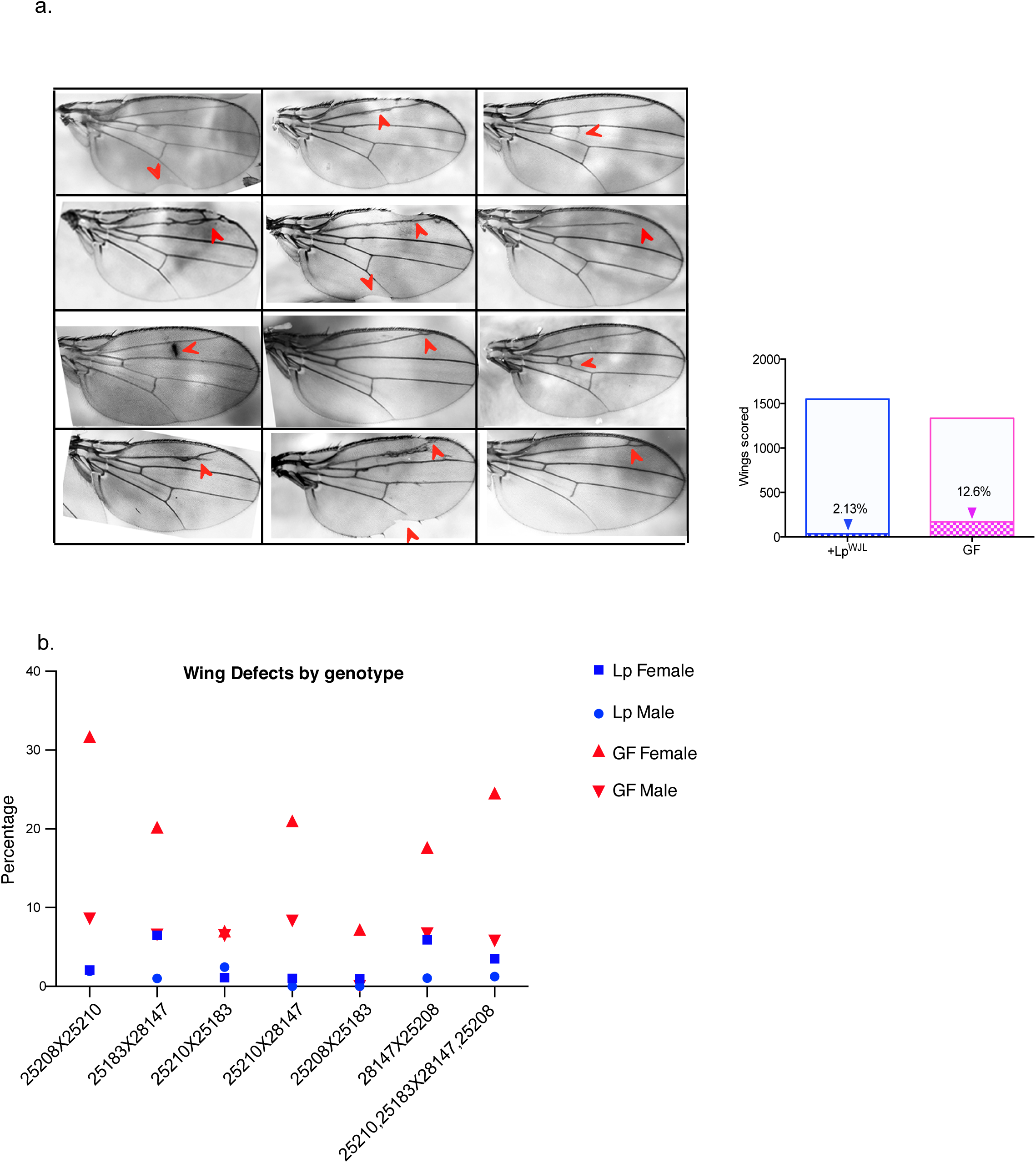
In the DGRP F_2_ progeny, *Lp*^*WJL*^ association provides robustness in wing developmental patterning. **a).**A compilation of representative images illustrating wing patterning anomalies in the GF DGRP F_2_ progeny, indicated by red arrows. The number of such patterning anomalies are compiled together for GF and *Lp*^*WJL*^ mono-associated flies (χ2 test, P<0.0001***, N_Lp_=1,551 N_GF_=1,335), and the percentage of defects are indicated inside each bar. **b).**The incidence of wing patterning defects separated by F_2_ genotypes. The Y-axis denotes the percentage of wings with aberrant patterning as represented in Figure **3a**.

### Compromising ROS activity impairs the buffering capacity of Lp^WJL^ without affecting bacteria growth

The wing anomalies in the GF F_2_ progeny highly resemble the phenotypes recently reported by *Santabarbara-Ruiz* et al, who blocked ROS activity through antioxidant feeding and induced regeneration defects in the wing[21]. We therefore repeated the DGRP F_2_ cross experiment with an additional condition by mixing the antioxidant molecule N-acetylcysteine (NAC) in the diet of mono-associated flies. NAC feeding did not compromise bacterial growth (Fig.S4b), but substantially diminished the buffering capacity of the bacteria (Fig.4). Specifically, variation in larval size (Fig.4a), developmental timing (Fig.4b and 4d) and adult emergence (Fig.4c and 4e) was significantly increased in NAC-fed larvae mono-associated with *Lp*^*WJL*^, to a level similar to or even higher than that in GF larvae. Wing patterning anomalies were also unmasked (Fig.4f). Therefore, blocking ROS activity through NAC-feeding suppresses the genetic buffering effect mediated by the gut bacteria. Jones *et al*. previously reported that acute exposure to *Lactobacillus plantarum* stimulates the *dNox*-dependent production of ROS in larval enterocytes, and subsequently increases the expression of genes involved in the Nrf2-mediated cyto-protection program[22, 23]. Future explorations are required to reconcile how ROS metabolism can be integrated into the molecular dialogue between the host and its intestinal microbiome to maintain robustness during development.

**Figure 4.**
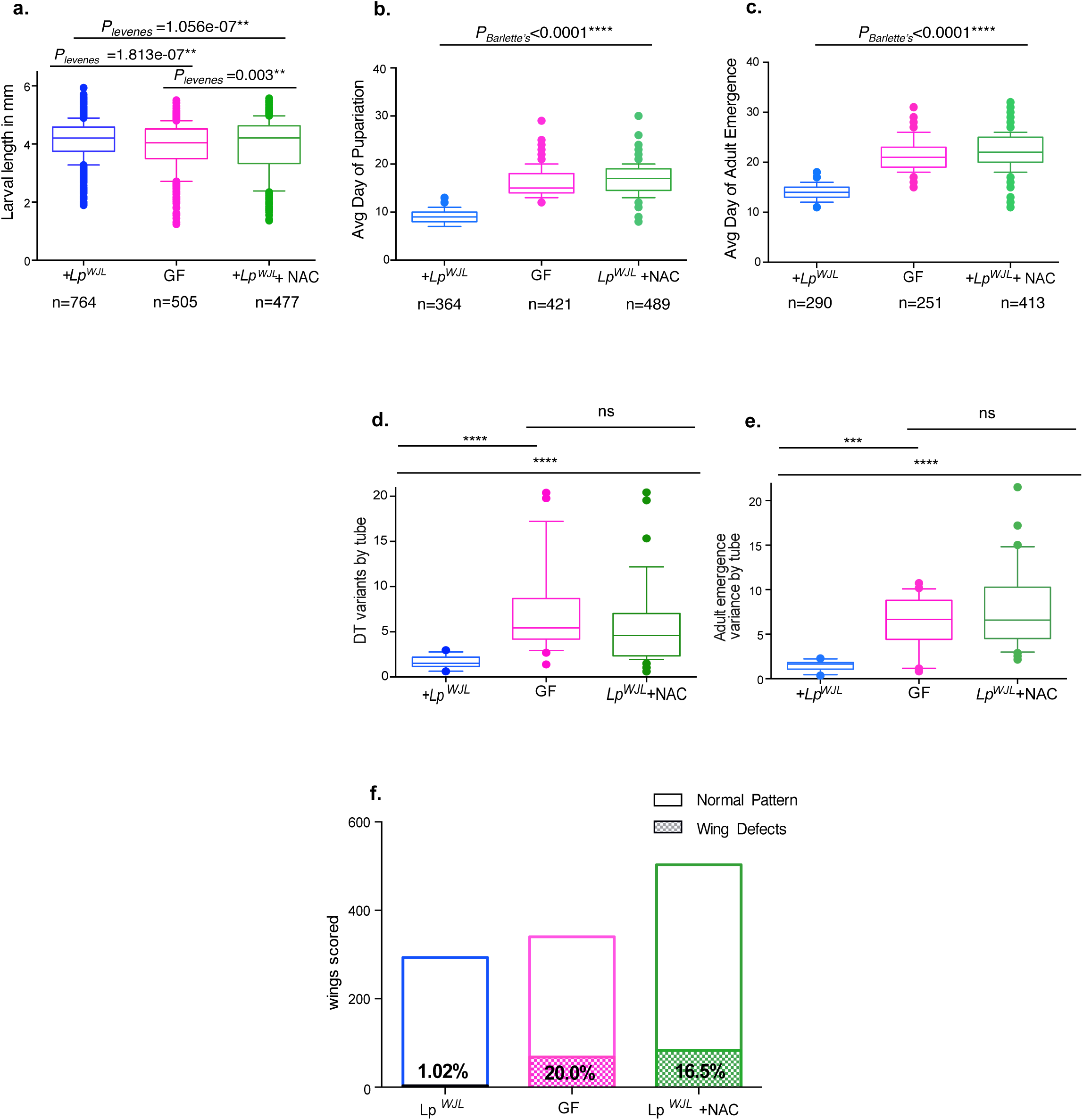
Blocking ROS activity by N-acetylcystein (NAC) compromises the *Lp*^*WJL*^ buffering capacity. **a).**In the DGRP F_2_ progeny, feeding *Lp*^*WJL*^ mono-associated animals with food supplemented with NAC increases the variances in size-matched larvae. Average Lp larval size: 4.08mm; average GF larval size: 3.83mm; average *Lp*^*WJL*^ +NAC larval size: 3.94mm. There is no size difference between GF and NAC treated flies associated with *Lp*^*WJL*^, p=0.064. CV_Lp_ =15.8%, CV_GF_= 20.8%; CV_Lp+NAC_=24.0%. **b).and c)**. NAC treatment to the Lp-associated animals also increases the variances of pupariation **(b)** and adult emergence **(c)**. The average day to become a pupa for *Lp*^*WJL*^ mono-associated larva: Day 8.9 (Var=2.13), for a GF larva: Day 16.1 (Var=8.27), for a NAC-treated, mono-associated larva: Day 16.8 (Var=8.36). The average day for an *Lp*^*WJL*^ mono-associated adult to emerge is: Day14.1 (Var=2.08), for a GF adult: Day 21 (Var= 8.3) and for a NAC-treated, mono-associated adult: Day 21.7 (Var=11.3). **d). and e)**. NAC treatment to the *Lp*^*WJL*^ mono-associated animals also increases the among-population variances of pupariation and adult emergence. Each data point represents the variance calculated based on the average day of pupariation (**d**) or adult emergence (**e**) from each tube housing approximately 40 animals. **f)**. Morphological defects in the wings are also significantly increased in NAC-treated mono-associated adults (D), (χ2 test, P<0.0001***) pink: GF (N=340); Blue: +*Lp*^*WJL*^ (N=293), Green: + *Lp*^*WJL*^ +NAC (N=503).

With a mono-association model, we unveiled that the *Drosophila* gut microbiota acts as a broad genetic buffer that safeguards the host’s genetic potential and confers developmental robustness when confronted with nutritional stress. This function may be a universal feature of beneficial microbes. In *Drosophila*, nutritional mutualism with commensals is facultative and volatile by nature[7, 24, 25]. Thus, the rapid acquisition or loss of particular gut community members can enable the developing host population to adjust its phenotypic range in response to the changing environment. The action of genetic buffering by microbiota in part invokes the concept of an “evolutionary capacitance”[2], and a future challenge is to prove if increased phenotypic variation due to loss of microbial buffering can be genetically assimilated in persistent nutritional stress. In line with our study, recent efforts by Elgart et al. showed that the effect of different mutant phenotypes is more pronounced in GF progeny than in their axenic parents[26], suggesting potential heritability of such variation. Lastly, by showing that the gut microbiota can mask the effect of cryptic genetic variation, our results may contribute to resolving the long-standing enigma of incomplete penetrance and expressivity in classical genetics and the “missing heritability” problem in contemporary genome-wide association studies.

## Acknowledgments

We would like to thank Benjamin Prud’homme and colleagues at ENS de Lyon for their critical reading of the manuscript and valuable suggestions, the Arthro-Tools platform the SFR Biosciences (UMS3444/US8) for the fly equipment and facility, the Bloomington Stock Centre and VDRC for fly lines. This work was supported by an international collaborative grant through the French “Agence Nationale de la Recherche” and the “Fond National Suisse pour la Recherche” (ANR-15-CE14-0028-01) awarded to F.L and B.D., an ERC starting grant (FP7/2007-2013-N_309704) awarded to F.L., a SystemsX.ch (AgingX) Grant awarded to B.D and Institutional Support by the EPFL (B.D.). CEI was funded by a Ph.D. fellowship from the Rhone-Alps region. F.L is supported by the FRM/FSER foundation, the FINOVI foundation and the EMBO Young Investigator Program. MBS, MF, ML, and B.D. were supported by AgingX (SystemsX.ch) and/or Institutional Support by the EPFL.

## Author contributions

D.M., M.B.S., B.D., and F.L. conceived the project and designed the experiments; D.M., and C.E.I., conducted all fly-related experiments; M.B.S., and M.L conducted the GWAS analysis; M.B.S, M.F. and V.B prepared the libraries and conducted single-larvae transcriptome analyses. P.J. conducted the multivariate statistical analyses; G.S., has identified the effect of NAC on *Lp*-mediated larval phenotypes. D.M., M.B.S., B.D., and F.L. analyzed the data. D.M. drafted the manuscript, D.M, M.B.S, B.D. and F.L revised the paper and wrote the final draft together.

## Methods

### Fly stocks and genetic crosses

Drosophila were kept at 25°C in a Panasonic Mir425 incubator with 12/12 hrs dark/light cycles. Routine stocks were kept on standard laboratory diet (see below “media preparation and NAC treatment”) The 53 DGRP lines were obtained from Bloomington Drosophila Stock Center.

Field-collected flies were trapped with rotten tomatoes in a garden in Solaize (France) and reared on a medium without chemical preservatives to minimize the modification to their gut microbiota[15]. One liter of media contains 15g inactivated yeast, 25g sucrose (Sigma Aldrich, ref. #84100), 80g cornmeal and 10g agar.

To generate DGRP F_2_s, four DGRP lines were selected for setting up seven different crosses: 25210 (RAL-859), 25183(RAL-335) are the lines with “large” larvae as germ-free, and 25208(RAL-820) and 28147(RAL-158) are the line with “small” larvae as germ-free (see figure legend Figure S3a).

All RNAi lines were crossed to the driver line *y,w;; tubulin*-GAL80^ts^, *daugtherless*- GAL4. To minimize lethality, we dampend the GAL4 strength by leaving the genetic crosses at 25°C. The following fly strains were used: *y,w*, UAS-*dpr-6*- IR(P{KK112634}VIE-260B), UAS-*CG13492*-IR, (*w*^*1118*^;P{GD14825}v29390), UAS-*daw*-IR(NIG #16987R-1), UAS-*sfl*-IR (*w*^*1118*^; P{GD2336}v5070), UAS-*arr*-IR (*w*^*1118*^; P{GD2617}v4818), UAS-*rg*-IR(*w*^*1118*^; P{GD8235}v17407), UAS-*bol*-IR(*w*^*1118*^;{GD10525}v21536), UAS-*glut1*-IR(*y*^*1*^ *v*^*1*^; P{TRiP.JF03060}attP2, Bloomington 28645), UAS-*CG32683*-IR (P{KK112515}VIE-260B), UAS-*CG42669*- IR(*w*^*1118*^;P{GD7292}v18081), UAS-*Eip75B*-IR (*w*^*1118*^; P{GD1434}v44851), UAS-*mCherry*-IR (*y*^*1*^ *v*^*1*^; P{CaryP}attP2), VDRC GD control (VDRC ID60000).

### GWAS and data computing of heritability indice

To calculate heritability, we estimated variance components using a random effects model using the lme4 R package[27]. Strain and experiment date were treated as random effects, and the heritability was calculated as VA/(VA+VD+VR), where VA is the additive genetic variance, and is equal to twice the Strain variance, VD is the experiment date variance, and VR is the residual variance. For the GWAS, we used the online tool specifically designed for the DGRPs (http://dgrp2.gnets.ncsu.edu/)[28, 29]. The Manhattan and QQ-plots were generated using R.

### Single larva transcriptome analysis

#### RNA extraction from single larvae

Larvae were handpicked under the microscope using forceps and transferred to Eppendorf tubes filled with 0.1 uL of beads and 350 uL of Trizol. The samples were then homogenized using a Precellys 24 Tissue Homogenizer at 6000 rpm for 30 seconds. After homogenization, the samples were transferred to liquid nitrogen for flash freezing and stored at –80°C. For RNA extraction, samples were thawed on ice, 350 uL of 100% Ethanol was then added to each sample before homogenizing again with the same parameters. Direct-zol(tm) RNA Miniprep R2056 Kit was used to extract RNA with these modifications: DNAse I treatment was skipped; after the RNA Wash step, an extra 2 min centrifugation step was added to remove residue wash. Lastly, the sample was eluted in 10 uL of water, incubated at room temperature for 2 min and then spun for 2 min to collect RNA. RNA was transferred to a low-binding 96 well plate and stored at −70°C.

#### RNA-sequencing

We prepared the libraries using the BRB-seq protocol and sequenced them using an Illumina NextSeq 500[11]. Reads from the BRB-seq protocol generates two fastq files: R1 containing barcodes and UMIs and R2 containing the read sequences. R2 fastq file was first trimmed for removing BRB-seq-specific adapter and polyA sequences using the BRB-seqTools v1.0 suite (available at http://github.com/DeplanckeLab/BRB-seqTools). We then aligned the trimmed reads to the Ensembl r78 gene annotation of the dm3 genome mixed with *the Lactobacillus Plantarum WJL* genome using STAR (Version 2.5.3a)[30], with default parameters (and extra “--outFilterMultimapNmax 1” parameter for completely removing multiple mapped reads). Then, using the BRB-seqTools v1.0 suite (available at http://github.com/DeplanckeLab/BRB-seqTools), we performed simultaneously the sample demultiplexing, and the count of reads per gene from the R1 FASTQ and the aligned R2 BAM files. This generated the count matrix that was used for further analyses. Genes were retained in the analysis if they had more than 10 reads in more than 50 samples. The data was subsequently transformed using the voom method. Differential expression was performed using the R Limma package[31, 32]. Genes with a log_2_ fold change greater than 2 and a Benjamini-Hochberg adjusted P-value less than 0.05 were considered differentially expressed. Since the library preparation was performed in two plates, hence introducing a batch effect, we used the duplicateCorrelation function and included the batch as a blocking variable. Prior to PCA analysis and standard deviation calculations, we removed the batch effect using the removeBatchEffects function and then used the princomp function. We used the cluster profiler package to perform GSEA analyses. The gmt file containing the gene ontology annotations was obtained from GO2MSIG data. Specifically, we used the high quality GO annotations for *Drosophila melanogaster*. For each GSEA analysis, we used 100,000 permutations to obtain adjusted p-values and only included gene set sizes to between 6 and 1000 genes. The raw expression data has been deposited in ArrayExpress (accession number: E-MTAB-6518)

#### RNA-sequencing

We prepared the libraries using the BRB-seq protocol and sequenced them using an Illumina NextSeq 500[11]. Reads from the BRB-seq protocol generates two fastq files: R1 containing barcodes and UMIs and R2 containing the read sequences. R2 fastq file was first trimmed for removing BRB-seq-specific adapter and polyA sequences using the BRB-seqTools v1.0 suite (available at http://github.com/DeplanckeLab/BRB-seqTools). We then aligned the trimmed reads to the Ensembl r78 gene annotation of the dm3 genome mixed with *the Lactobacillus Plantarum WJL* genome using STAR (Version 2.5.3a)[30], with default parameters (and extra “--outFilterMultimapNmax 1” parameter for completely removing multiple mapped reads). Then, using the BRB-seqTools v1.0 suite (available at http://github.com/DeplanckeLab/BRB-seqTools), we performed simultaneously the sample demultiplexing, and the count of reads per gene from the R1 FASTQ and the aligned R2 BAM files. This generated the count matrix that was used for further analyses. The data was subsequently transformed using the voom method and analyzed using the R Limma package[31, 32].

The raw expression data of BRB-Seq has been deposited in ArrayExpress (accession number: E-MTAB-6518)

### The making and maintenance of germ-free flies

Axenic flies were generated by dechorionating embryos with 50% household bleach for five minutes; eggs were then washed in successive 70% ethanol and sterile distilled water for three minutes each. After washing, eggs were transferred to tubes containing standard diet and a cocktail of antibiotics containing 50µg/mL ampicillin, 50µg/mL kanamycin, 15µg/mL erythromycin, 50µg/mL tetracyclin for stock maintenance. Axeny was routinely verified by plating larvae and adult lysates on LB and MRS plates. For experiments food without antibiotics was used.

### Media preparation and NAC treatment

Standard laboratory fly food consists of 50g/L inactivated yeast (Springaline(tm)), 80g/L cornmeal, 7.14g/L agar, 5.12g/L Moldex (Sigma M-50109) and 0.4% propionic acid. Where applicable, experiments comparing variations in larval size, developmental timing, adult emergence were performed on diet with 6g or 8g inactivated yeast per liter of media while keeping the same concentrations for the other ingredients. Where appropriate, 1.7g/L of N-Acetylcystein (SigmaA7250-25g) was added to the low-protein diet.

### Larval Length Measurement

All live *Drosophila* larvae were collected from each nutritive cap containing low yeast diet by temporary immersion in sterile PBS, transferred on a microscopy slide, killed with a short pulse of heat (5 sec at 90°C), mounted with 80% glycerol/PBS. The images were taken with the Leica stereomicroscope M205FA and the lengths of individual larvae were measured using ImageJ software[33]. For each DGRP strain and each cross and/or condition, at least three biological replicates were generated.

### Developmental timing and Adult emergence

Developmental timing and adult emergence of the flies were quantified by counting the number of individuals appearing every 24 hours until the last pupa/adult emerges. Each animal is assigned to the number that corresponds to the day it appeared, and the population mean and variance were calculated based on the cumulative numbers.

### Adult trait measurements

2-3 days old adult flies were anesthetized with CO_2_ and immersed in 70% ethanol, and the individual body and its corresponding organ (wing and eye) were imaged under a Leica M205 stereomicroscope. Specifically, the adult body length was measured from the top of the head to the tip of the abdomen. The eye area was measured by manually tracing the circumference of both eyes. The wings were gently nipped at the base of the hinge and imaged, and the area was measured by tracing the edge of the wing. All images were taken measured using ImageJ software

### Bacteria culture and mono-association

For each mono-association experiment, Lp^*WJL*^ [34] was grown in Man, Rogosa and Sharpe (MRS) medium (Difco, ref. #288110) over-night at 37°C, and diluted to O.D.=0.5 the next morning to inoculate 40 freshly laid eggs on a 55mm petri dish or standard 28mm tubes containing fly food of low yeast content. The inoculum corresponds to about 5×10^7^ CFUs. Equal volume of sterile PBS was spread on control axenic eggs.

To contaminate the garden-collected flies with their own microbiota, eggs were dechorionated and directly seeded onto appropriate food caps. Sterile PBS was used to wash the side of the bottles where the adult wild flies were raised to recover more fecal content, and 300 ul of the wash was inoculated to the dechorionated eggs. For GF control, 300 ul of sterile PBS was used to inoculate the dechorionated eggs. The microbial composition of this microbiota can be founded here[15].

### Bacteria niche load

Five to six 24 hour old germ-free larvae were collected from the low-protein diet food cap and transferred to a microtube containing 400ul of low-protein diet, and inoculated with 50ul of *Lp*^*WJL*^ of 0.5 O.D.. On the day of harvest, ∼0.75-1mm glass micro-beads and 900µl PBS were added to each microtube and the entire content of the tube was homogenized with the Precellys-24 tissue homogenizer (Bertin Technologies). Lysate dilutions (in PBS) are plated on MRS agar with Easyspiral automatic plater (Intersciences). The MRS agar plates were incubated for 24h at 37°C. The CFU/ml count was calculated based on the readings by the automatic colony counter Scan1200 (Intersciences)

### Statistical Analysis and data representation

GraphPad Prism software version 6.0f for Macintosh (GraphPad Software, La Jolla California USA, www.graphpad.com) was used to compare GF and *Lp*^*WJL*^-associated conditions for larval length, developmental timing, adult emergence, allometry and linear regression analysis for the buffering effect. For small samples with less than 10 data points, nonparametric analysis was conducted. R-studio was used to conduct Levene’s test and multivariate analyses. For all experiments, the p-values were reported on the corresponding figure panels only when inferior to 0.05.

---

**Figure S1.**
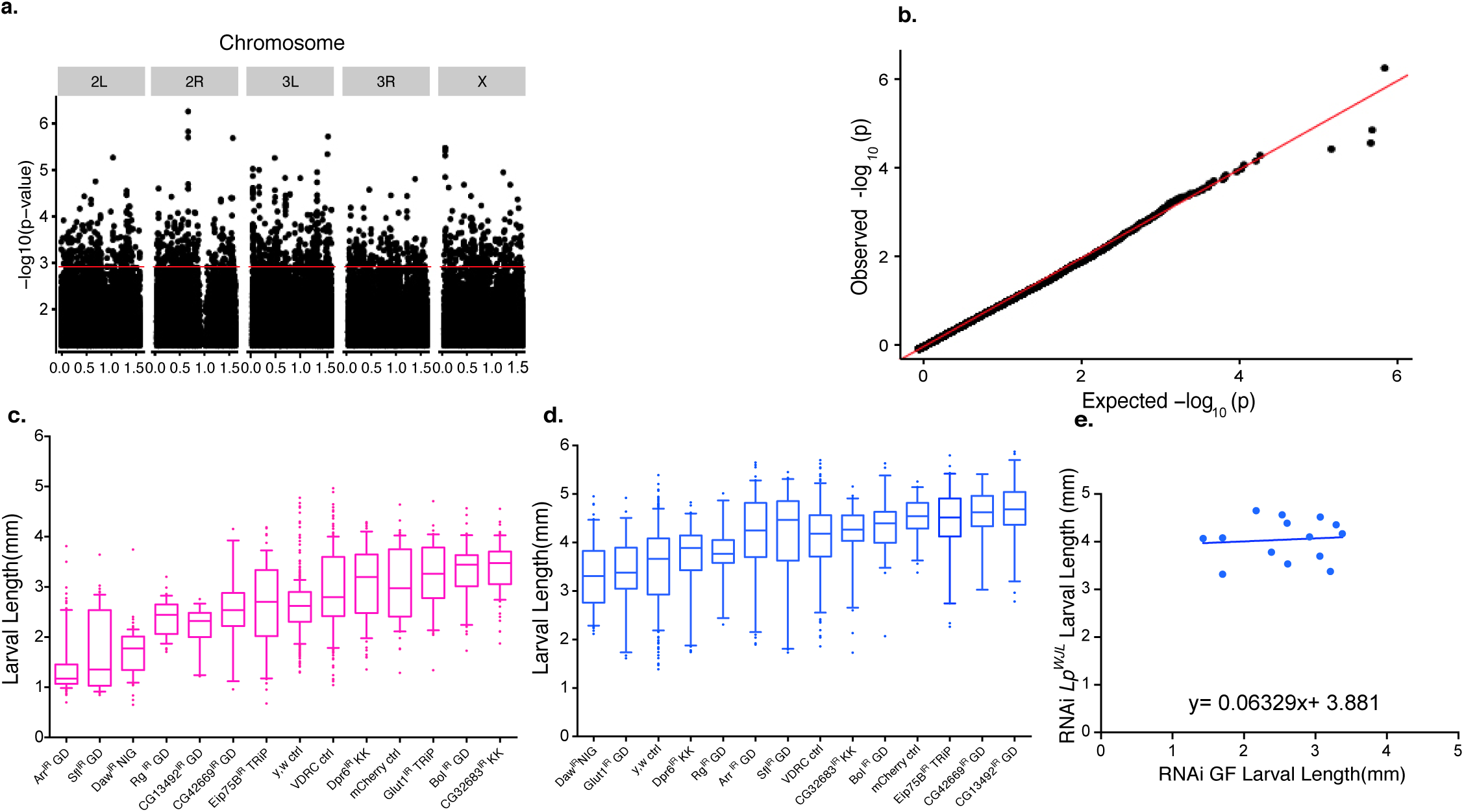
**a)**. Manhattan plot of the GWAS performed on the average larval length fold change per DGRP line. We used the DGRP2 website for the association analysis (http://dgrp2.gnets.ncsu.edu/)[28, 29]. **b)**. Quantile-Quantile plot of the GWAS results. **c). and d)**. Box and whiskers plots illustrating the effect of RNAi knockdown on larval length on day 7 AEL. Each bar represents the average length from pooled 3-5 biological replicates from either condition, with 15-40 larvae in each replicate. **c:** GF. **d:** *Lp*^*WJL*^. Three different control knockdowns were used: one control fly strain recommended by VDRC for RNAi constructs obtained from VDRC, one control strain (against mCherry) recommended by the Harvard TRiP collection, and the *y,w* strain from Bloomington. All control and RNAi strains were crossed to *y,w;; tubulin*-GAL80^ts^, *daugtherless*-GAL4. “GD” refers to the VDRC RNAi GD collection. “KK” refers to the VDRC RNAi KK collection. For specific genotypes, refer to Material and Methods. **e)**. *Lp*^*WJL*^ also buffers growth differences in various RNAi knock-down experiments for each of the candidate genes. Each data point represents the intercept of the GF length and its corresponding mono-associated length at Day 7 for the RNAi knockdown experiment. (Null hypothesis: Slope =1. *P=0.0008*, the null hypothesis is therefore rejected). These data points were fitted into an unconstraint model. For specific genotypes, we refer to Table 2 and Methods.

**Figure S2.**
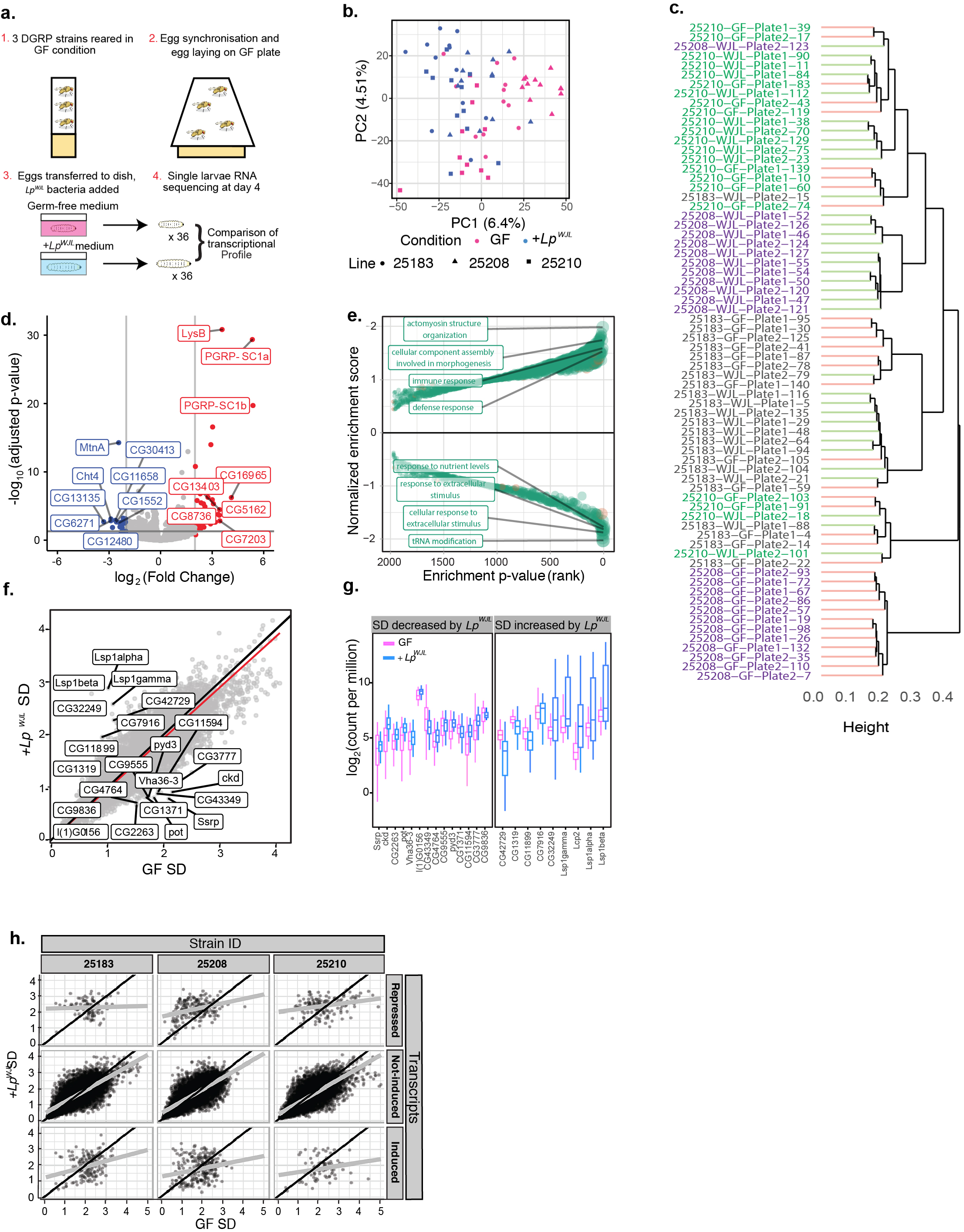
**a)**. Experimental setup to perform BRB-seq-based transcriptomics on individual larvae. Flies from three DGRP strains were reared in GF conditions. Egg-laying was synchronized and embryos were transferred to food caps: three left germ-free (1X PBS) and three inoculated with *Lp*^*WJL*^ (OD 0.5 in 1x PBS). At day 4, single larvae were collected from all plates, RNA extraction and RNA sequencing were performed. In sum, 12 larvae were collected per line for each condition, totaling 72 single larval transcriptomes. **b)**. Principal component plot of the corrected expression data after batch correction. **c)**. Hierarchical clustering of the transcriptomic data using the Ward’s method. A batch effect of plate was corrected prior to clustering. The genotypes are color coded (Green: 25210, violet: 25208, black: 25183). The red “branches” of the cluster represent GF samples, and green ones represent mono-associated samples. **d)**. The observed effect of *Lp*^*WJL*^ mono-association on gene expression is consistent with our previous findings, thus validating our transcriptome approach on individual larvae. The horizontal grey line represents the 0.05 FDR-corrected p-value threshold. The vertical lines are the −2 and 2 log2 (Fold Change) thresholds. Genes in red, such as *LysB, PGRP-SC1a&b* are significantly up-regulated; they are predominantly involved in host immune and defense response (see also S1e); genes in blue are significantly down-regulated. Several representative genes of the top differentially regulated genes from each category are highlighted. **e)**. Gene set enrichment analysis on biological process gene ontology (GO) terms based on the effect of *Lp*^*WJL*^ mono-association. Gene sets in orange were derived from GLAD[35], whereas green gene sets were extracted from GO2MSIG[36]. Note that “immune response”, “defense response” and “cellular component assembly involved in morphogenesis” are among the most up-regulated gene sets by mono-association (top panel), and genes associated to “response to nutrient levels”, “cellular response to starvation” and “t-RNA modification” were down-regulated by *Lp*^*WJL*^ (bottom panel). Therefore, both microbe detection and nutrient adaptation drive the most significantly detected transcriptomic changes in mono-associated larvae. **f)**. Scatterplot of the standard deviation in expression level of each gene in the GF and *Lp*^*WJL*^ mono-associated condition. The black line represents the theoretical slope of 1 and intercept 0. The red line is a linear fit of the points. Labelled genes show the highest relative change in their standard deviation, as determined by the absolute value of log_2_(SD_*LpWJL*_/SD_GF_). **g)**. Box and whiskers plots showing the expression levels of genes with high relative change in standard deviation, regardless whether the genes themselves were up- or down-regulated. Among the genes whose expression variation decreased the most upon *Lp*^*WJL*^ association are *Ssrp*, a member of the FACT chromatin complex[37, 38], and many cuticle-related proteins (left panel), whereas for genes induced by *Lp*^*WJL*^, such as <u>L</u>arval <u>s</u>erum <u>p</u>roteins (Lsp1s), more expression variation is detected (right panel). **h)**. Scatterplots of standard deviations of each gene calculated by genotype. Genes were faceted by how their differential expression alters within each strain in both GF and *Lp*^*WJL*^ mono-associated conditions: repressed (top panel), non-induced (middle panel) and induced (bottom panel). The black lines represent the theoretical slope of 1 and intercepts 0, the grey lines are the linear fit to the data. Since transcripts specifically modulated by *Lp*^*WJL*^ tend to have incomparable SD, we assessed GO enrichment only on non-differentially expressed genes (**see Fig.1j**)

**Figure S3.**
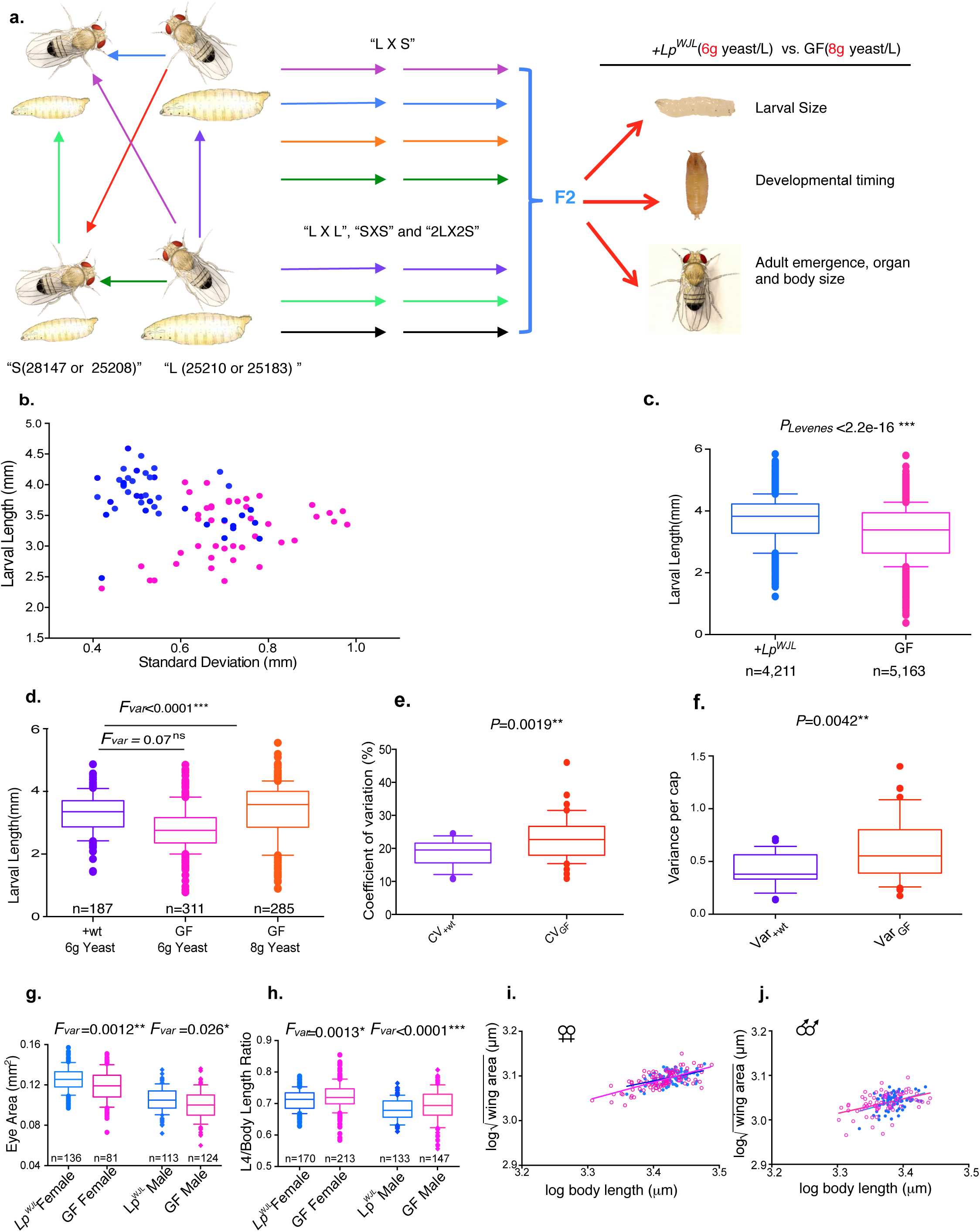
**a)**. A diagram illustrating DGRP crosses to generate the F_2_ generation for studying variation in larval size, pupariation and adult emergence. 25210 (RAL-859), 25183(RAL-335) are the lines with the “large” (“L”) larvae as germ-free, and 25208(RAL-820) and 28147(RAL-158) are the lines with the “small” larvae as germ-free (“S”). Seven possible crosses are set up: 25210X25183 (“LXL”), 25208X28147(“SXS”), 25210X25208, 25183X25208, 25210X28147, 25183X28147 are the four “LXS” crosses, and 25183 and 25210 × 25208 and 28147 is the “2L X 2S” cross. **b)**. A scatter plot showing how standard deviation (SD) changes as a function of larval length, and how such change differs in the DGRP F_2_ GF (pink) and *Lp*^*WJL*^ mono-associated (blue) populations (see also Figure 2a and Methods for detailed schemes). Each data point represents the intercept of an SD value and its corresponding average larval length in a particular cross. Each SD and average length was derived from larvae measurements gathered from at least 3 biological replicates from either GF or *Lp*^*WJL*^ mono-associated conditions. Each replicate contains 10-40 larvae. **c)**. Box and Whisker graph illustrating the average length and standard deviation from pooled GF (pink) and *Lp*^*WJL*^ mono-associated DGRP (blue) F2 larvae, pooled from all the crosses in all three different repeats (Average GF larval length: 3.29mm; average Lp mono-associated larval length: 3.71mm; CV_GF_=24.9%, CV_Lp_=19.5%). **d)**. One representative experiment showing that re-associating the field-collected flies tends to buffer the variability in body length in size-matched larvae. The purple box represents body length from wild larvae grown on media contaminated with their untreated parents’ fecal matter. Average GF larval length grown on 6g/L yeast media: 2.81mm; average GF larval length grown on 8g/L yeast media: 3.36mm: average re-associated larval length (“+wt”): 3.07 mm; P= 0.338. CV_GF_ (6g/L, pink)= 24.9%, CV_GF_ (8g/L, orange)= 27.0%, CVwt (purple)= 18.9%. **e)**. **and f)**. The compiled CV values (**e**.) and variances (**f**.) derived from each low-yeast cap containing 40∼50 field-collected larvae. The average CV and variance are lower in the population re-associated with its own microbiota (purple) than in the GF population (orange) **g)**. In both male (lozenge) and female (circle) adults, the variances in eye size are greater in GF F_2_ progeny. The difference in mean eye area, for females P<0.0001***; for males, P=0.0013**. **h)**. The length of the L4 vein in the wing is used as a proxy of the wing length. In the accumulated ratios of wing length over body length, the variances are greater in the GF flies (The difference in average L4/ body length, for females P<0.0028**; for males, P=0.02*). **i) and j)**. Scatter plots illustrating the allometric relationship between wing area and body size in female (i) and male (j) DGRP F_2_ adults. Pink open circles: GF, blue filled circles: *Lp*^*WJL*^. Each line represents the allometric slope of the data points shown by the same color. Either in males or females, there is no difference in allometric slope between the GF and mono-associated population. For GF females, Y_GF_ = 0.3963*X + 1.738, 95%C.I.= 0.3117 to 0.4810; for *Lp*^*WJL*^ females, Y_Lp_ = 0.2978*X + 2.076, 95%C.I.=0,1785 to 0,4172, P=0.203, n.s; for GF males, Y_GF_ = 0.3261*X + 1.939, 95%C.I.=0.1725 to 0.4796; for *Lp*^*WJL*^ males, Y_Lp_= 0.4141*X + 1.639, 95% C.I. =0.1842 to 0.6439, P=0.55, ns.

**Figure S4.**
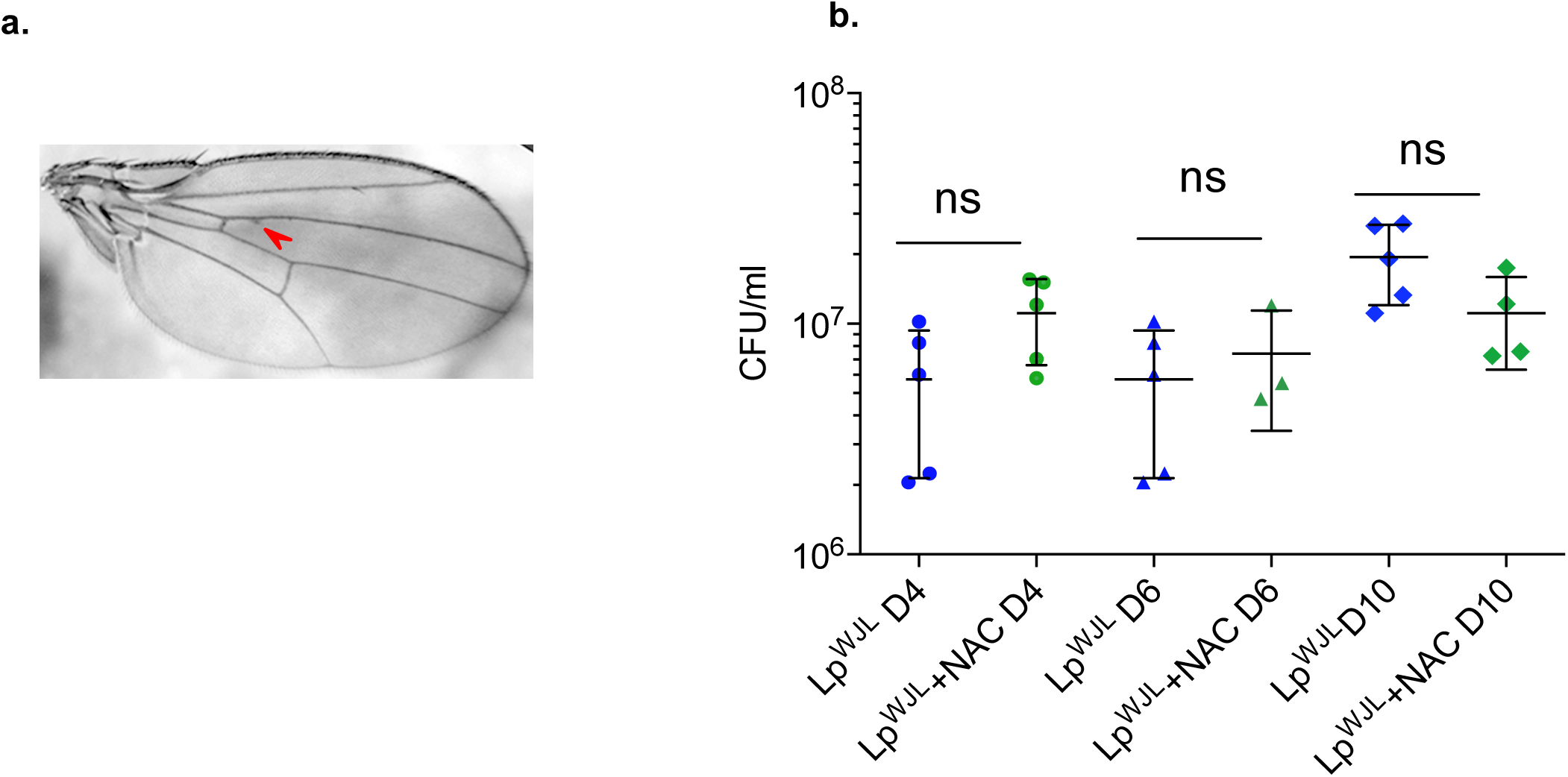
**a)**. An image of a wing of an *Lp*^*WJL*^ adult is shown, as a representation of the most visible “defect” ever observed in mono-associated adults. Red arrow points to the subtle vein tissue thickening. We included these as “defects” in the *Lp*^*WJL*^ F_2_ population in the analyses presented in Figure 3a, 3b, and 4f. **b)**. Bacterial niche load (NL) evolution (“Niche” is defined as the substrate with both larvae and bacteria present) during the course of larval development with *Lp*^*WJL*^ with or without NAC treatment (Day 4, Day 6 and Day 10).

**Table S1.** Average D7 larvae length for individual DGRP lines (Related to Figure1) *GF: germ-free *\*Lp*^*WJL*^: Lactobacillus plantarum, stain name: WJL *SD: standard deviation

| **DGRP**<br>**Lines** | **GF\***<br>**Length(mm)** | **GF SD\*(mm)** | **$Lp^{WJL*}$**<br>**Length(mm)** | **$Lp^{WJL}$**<br>**SD(mm)** | **$Lp^{WJL}$**<br>**/GF**<br>**Ratio** |
| --- | --- | --- | --- | --- | --- |
| 25174 | 2.193 | 0.584 | 3.637 | 0.895 | 1.658 |
| 25175 | 2.693 | 0.687 | 4.496 | 0.659 | 1.670 |
| 25176 | 1.443 | 0.536 | 3.903 | 0.648 | 2.704 |
| 25180 | 2.151 | 0.454 | 3.795 | 0.635 | 1.764 |
| 25181 | 2.374 | 0.824 | 4.224 | 0.946 | 1.779 |
| 25182 | 2.108 | 0.451 | 3.293 | 0.859 | 1.562 |
| 25183 | 2.961 | 0.657 | 4.066 | 0.548 | 1.373 |
| 25184 | 1.957 | 0.53 | 4.323 | 0.587 | 2.209 |
| 25185 | 2.459 | 0.681 | 3.93 | 0.722 | 1.598 |
| 25186 | 2.278 | 0.667 | 4.289 | 0.803 | 1.883 |
| 25187 | 2.109 | 0.479 | 3.798 | 0.744 | 1.801 |
| 25188 | 2.253 | 0.421 | 4.202 | 0.786 | 1.865 |
| 25189 | 2.586 | 0.393 | 3.448 | 0.876 | 1.333 |
| 25190 | 2.292 | 0.512 | 3.976 | 0.941 | 1.735 |
| 25191 | 2.348 | 0.428 | 3.953 | 0.797 | 1.684 |
| 25192 | 2.194 | 0.401 | 4.145 | 0.731 | 1.889 |
| 25193 | 2.414 | 0.582 | 4.05 | 0.782 | 1.678 |
| 25194 | 2.506 | 0.558 | 4.195 | 0.508 | 1.674 |
| 25195 | 2.07 | 0.402 | 3.635 | 0.867 | 1.756 |
| 25197 | 1.944 | 0.397 | 3.73 | 0.734 | 1.919 |
| 25198 | 2.051 | 0.394 | 3.936 | 0.673 | 1.919 |
| 25199 | 1.514 | 0.524 | 3.78 | 0.753 | 2.497 |
| 25200 | 2.869 | 0.752 | 4.227 | 0.605 | 1.473 |
| 25201 | 2.182 | 0.347 | 4.186 | 0.601 | 1.918 |
| 25202 | 2.273 | 0.639 | 3.85 | 0.792 | 1.694 |
| 25203 | 1.541 | 0.513 | 4.158 | 0.755 | 2.698 |
| 25204 | 1.686 | 0.678 | 4.088 | 0.774 | 2.425 |
| 25205 | 2.351 | 0.567 | 3.77 | 0.606 | 1.604 |
| 25206 | 2.5 | 0.643 | 4.173 | 0.619 | 1.669 |
| 25207 | 2.028 | 0.481 | 3.896 | 0.811 | 1.921 |
| 25208 | 1.649 | 0.443 | 4.103 | 0.947 | 2.488 |
| 25209 | 2.187 | 0.67 | 4.232 | 0.819 | 1.935 |
| 25210 | 2.772 | 0.633 | 4.03 | 0.466 | 1.454 |
| 25445 | 2.01 | 0.468 | 3.956 | 0.668 | 1.968 |
| 25744 | 2.097 | 0.34 | 4.235 | 0.666 | 2.020 |
| 25745 | 2.501 | 0.612 | 4.051 | 0.599 | 1.620 |
| 28132 | 2.828 | 0.684 | 4.485 | 0.534 | 1.586 |
| 28134 | 1.854 | 0.383 | 4.144 | 0.479 | 2.235 |
| 28136 | 1.707 | 0.415 | 4.204 | 0.548 | 2.463 |
| 28138 | 1.38 | 0.487 | 4.318 | 0.693 | 3.129 |
| 28142 | 2.938 | 0.836 | 4.487 | 0.489 | 1.527 |
| 28146 | 2.077 | 0.36 | 4.564 | 0.915 | 2.197 |
| 28147 | 1.575 | 0.552 | 4.061 | 0.728 | 2.578 |
| 28153 | 2.298 | 0.329 | 3.97 | 0.541 | 1.728 |
| 28154 | 2.256 | 0.339 | 4.365 | 0.482 | 1.935 |
| 28160 | 2.51 | 0.662 | 4.118 | 0.714 | 1.640 |
| 28164 | 2.394 | 0.448 | 4.207 | 0.584 | 1.757 |
| 28166 | 2.163 | 0.402 | 4.489 | 0.642 | 2.075 |
| 28173 | 2.039 | 0.309 | 4.122 | 0.697 | 2.022 |
| 28192 | 2.141 | 0.506 | 4.286 | 0.659 | 2.002 |
| 28194 | 2.269 | 0.565 | 4.424 | 0.72 | 1.950 |
| 28197 | 2.89 | 0.742 | 4.547 | 0.519 | 1.573 |
| 28208 | 2.339 | 0.438 | 4.14 | 0.705 | 1.767 |

\*GF: germ-free
\*SD: standard deviation

**Table S2.**
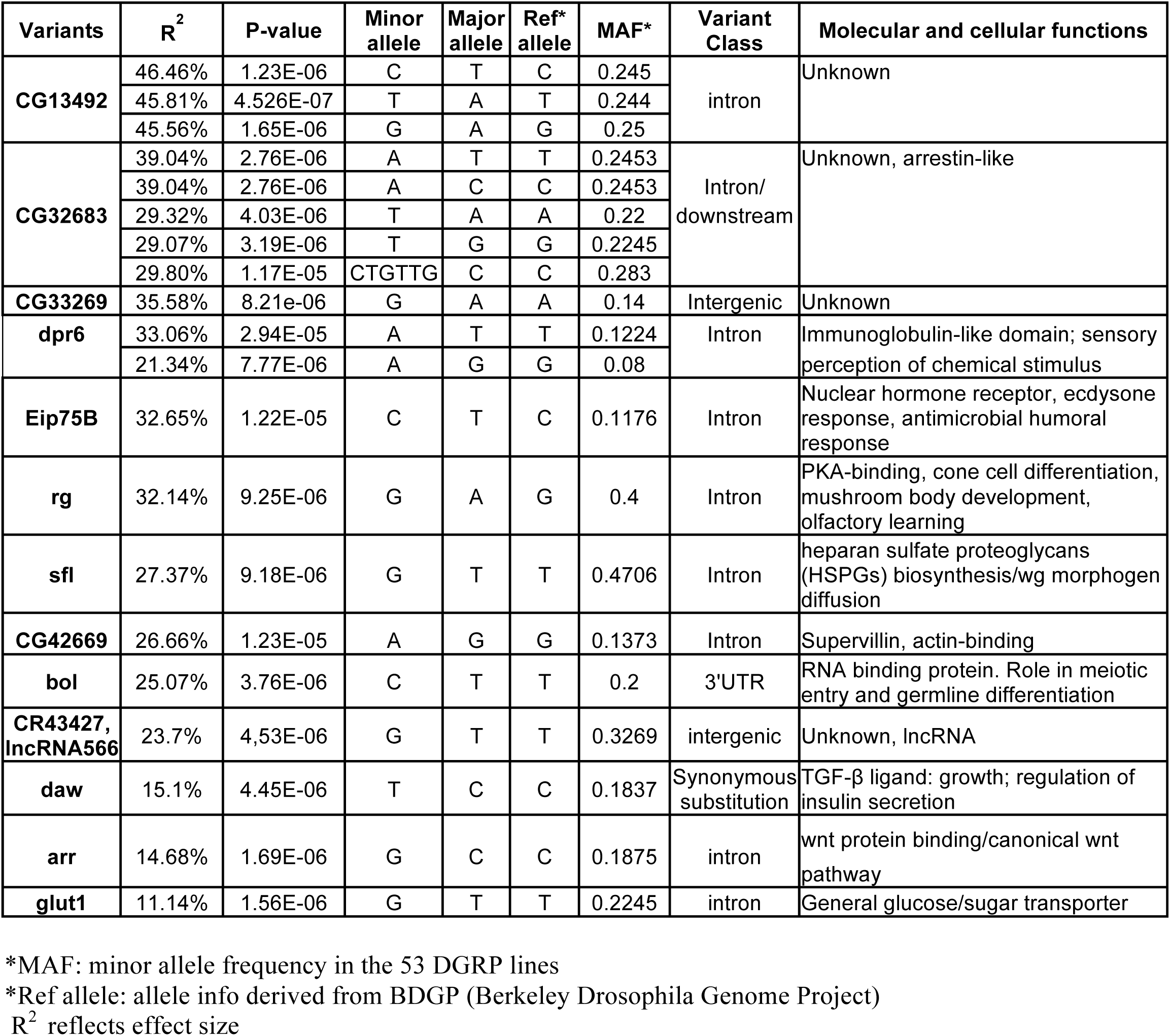
Variants associated with the growth benefits conferred by *Lactobacillus plantarum* (*Lp*^*WJL*^) (Related to Figure S1)

| Variants | R<sup>2</sup> | P-value | Minor allele | Major allele | Ref* allele | MAF* | Variant Class | Molecular and cellular functions |
| --- | --- | --- | --- | --- | --- | --- | --- | --- |
| **CG13492** | 46.46% | 1.23E-06 | C | T | C | 0.245 | intron | Unknown |
|  | 45.81% | 4.526E-07 | T | A | T | 0.244 |  |  |
|  | 45.56% | 1.65E-06 | G | A | G | 0.25 |  |  |
| **CG32683** | 39.04% | 2.76E-06 | A | T | T | 0.2453 | Intron/<br>downstream | Unknown, arrestin-like |
|  | 39.04% | 2.76E-06 | A | C | C | 0.2453 |  |  |
|  | 29.32% | 4.03E-06 | T | A | A | 0.22 |  |  |
|  | 29.07% | 3.19E-06 | T | G | G | 0.2245 |  |  |
|  | 29.80% | 1.17E-05 | CTGTTG | C | C | 0.283 |  |  |
| **CG33269** | 35.58% | 8.21e-06 | G | A | A | 0.14 | Intergenic | Unknown |
| **dpr6** | 33.06% | 2.94E-05 | A | T | T | 0.1224 | Intron | Immunoglobulin-like domain; sensory perception of chemical stimulus |
|  | 21.34% | 7.77E-06 | A | G | G | 0.08 |  |  |
| **Eip75B** | 32.65% | 1.22E-05 | C | T | C | 0.1176 | Intron | Nuclear hormone receptor, ecdysone response, antimicrobial humoral response |
| **rg** | 32.14% | 9.25E-06 | G | A | G | 0.4 | Intron | PKA-binding, cone cell differentiation, mushroom body development, olfactory learning |
| **sfl** | 27.37% | 9.18E-06 | G | T | T | 0.4706 | Intron | heparan sulfate proteoglycans (HSPGs) biosynthesis/wg morphogen diffusion |
| **CG42669** | 26.66% | 1.23E-05 | A | G | G | 0.1373 | Intron | Supervillin, actin-binding |
| **bol** | 25.07% | 3.76E-06 | C | T | T | 0.2 | 3'UTR | RNA binding protein. Role in meiotic entry and germline differentiation |
| **CR43427, lncRNA566** | 23.7% | 4,53E-06 | G | T | T | 0.3269 | intergenic | Unknown, lncRNA |
| **daw** | 15.1% | 4.45E-06 | T | C | C | 0.1837 | Synonymous substitution | TGF-β ligand: growth; regulation of insulin secretion |
| **arr** | 14.68% | 1.69E-06 | G | C | C | 0.1875 | intron | wnt protein binding/canonical wnt pathway |
| **glut1** | 11.14% | 1.56E-06 | G | T | T | 0.2245 | intron | General glucose/sugar transporter |

\*MAF: minor allele frequency in the 53 DGRP lines
\*Ref allele: allele info derived from BDGP (Berkeley Drosophila Genome Project)
R<sup>2</sup> reflects effect size

**Table S3.**
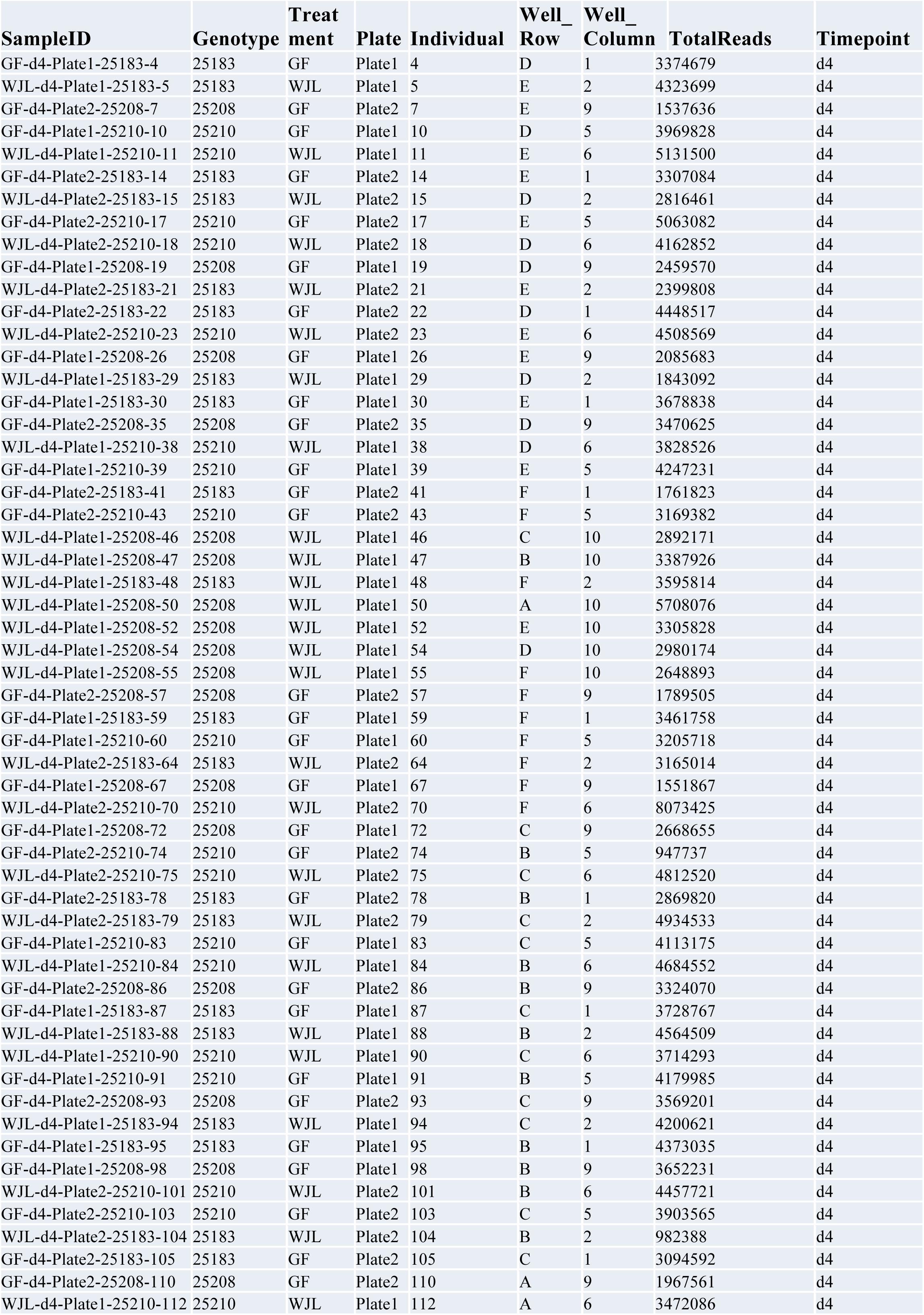
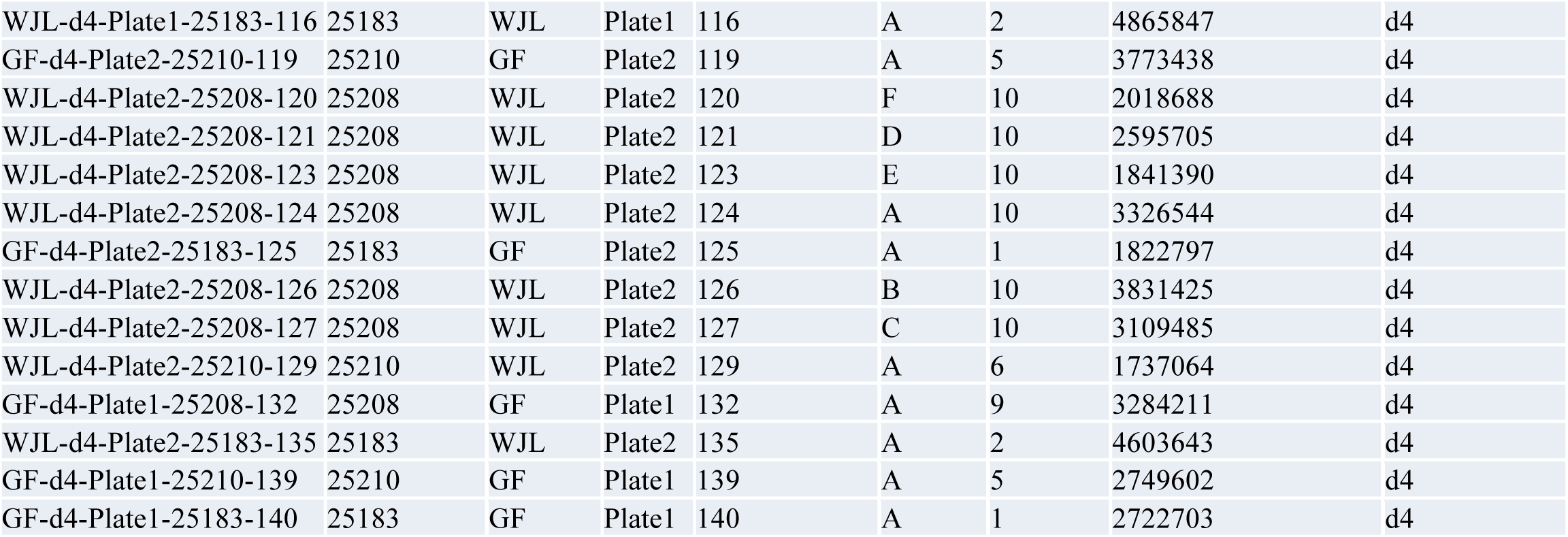
Individual larval transcriptome sample list (Related to Figure S2)

| SampleID | Genotype | Treatment | Plate | Individual | Well_<br>Row | Well_<br>Column | TotalReads | Timepoint |
| --- | --- | --- | --- | --- | --- | --- | --- | --- |
| GF-d4-Plate1-25183-4 | 25183 | GF | Plate1 | 4 | D | 1 | 3374679 | d4 |
| WJL-d4-Plate1-25183-5 | 25183 | WJL | Plate1 | 5 | E | 2 | 4323699 | d4 |
| GF-d4-Plate2-25208-7 | 25208 | GF | Plate2 | 7 | E | 9 | 1537636 | d4 |
| GF-d4-Plate1-25210-10 | 25210 | GF | Plate1 | 10 | D | 5 | 3969828 | d4 |
| WJL-d4-Plate1-25210-11 | 25210 | WJL | Plate1 | 11 | E | 6 | 5131500 | d4 |
| GF-d4-Plate2-25183-14 | 25183 | GF | Plate2 | 14 | E | 1 | 3307084 | d4 |
| WJL-d4-Plate2-25183-15 | 25183 | WJL | Plate2 | 15 | D | 2 | 2816461 | d4 |
| GF-d4-Plate2-25210-17 | 25210 | GF | Plate2 | 17 | E | 5 | 5063082 | d4 |
| WJL-d4-Plate2-25210-18 | 25210 | WJL | Plate2 | 18 | D | 6 | 4162852 | d4 |
| GF-d4-Plate1-25208-19 | 25208 | GF | Plate1 | 19 | D | 9 | 2459570 | d4 |
| WJL-d4-Plate2-25183-21 | 25183 | WJL | Plate2 | 21 | E | 2 | 2399808 | d4 |
| GF-d4-Plate2-25183-22 | 25183 | GF | Plate2 | 22 | D | 1 | 4448517 | d4 |
| WJL-d4-Plate2-25210-23 | 25210 | WJL | Plate2 | 23 | E | 6 | 4508569 | d4 |
| GF-d4-Plate1-25208-26 | 25208 | GF | Plate1 | 26 | E | 9 | 2085683 | d4 |
| WJL-d4-Plate1-25183-29 | 25183 | WJL | Plate1 | 29 | D | 2 | 1843092 | d4 |
| GF-d4-Plate1-25183-30 | 25183 | GF | Plate1 | 30 | E | 1 | 3678838 | d4 |
| GF-d4-Plate2-25208-35 | 25208 | GF | Plate2 | 35 | D | 9 | 3470625 | d4 |
| WJL-d4-Plate1-25210-38 | 25210 | WJL | Plate1 | 38 | D | 6 | 3828526 | d4 |
| GF-d4-Plate1-25210-39 | 25210 | GF | Plate1 | 39 | E | 5 | 4247231 | d4 |
| GF-d4-Plate2-25183-41 | 25183 | GF | Plate2 | 41 | F | 1 | 1761823 | d4 |
| GF-d4-Plate2-25210-43 | 25210 | GF | Plate2 | 43 | F | 5 | 3169382 | d4 |
| WJL-d4-Plate1-25208-46 | 25208 | WJL | Plate1 | 46 | C | 10 | 2892171 | d4 |
| WJL-d4-Plate1-25208-47 | 25208 | WJL | Plate1 | 47 | B | 10 | 3387926 | d4 |
| WJL-d4-Plate1-25183-48 | 25183 | WJL | Plate1 | 48 | F | 2 | 3595814 | d4 |
| WJL-d4-Plate1-25208-50 | 25208 | WJL | Plate1 | 50 | A | 10 | 5708076 | d4 |
| WJL-d4-Plate1-25208-52 | 25208 | WJL | Plate1 | 52 | E | 10 | 3305828 | d4 |
| WJL-d4-Plate1-25208-54 | 25208 | WJL | Plate1 | 54 | D | 10 | 2980174 | d4 |
| WJL-d4-Plate1-25208-55 | 25208 | WJL | Plate1 | 55 | F | 10 | 2648893 | d4 |
| GF-d4-Plate2-25208-57 | 25208 | GF | Plate2 | 57 | F | 9 | 1789505 | d4 |
| GF-d4-Plate1-25183-59 | 25183 | GF | Plate1 | 59 | F | 1 | 3461758 | d4 |
| GF-d4-Plate1-25210-60 | 25210 | GF | Plate1 | 60 | F | 5 | 3205718 | d4 |
| WJL-d4-Plate2-25183-64 | 25183 | WJL | Plate2 | 64 | F | 2 | 3165014 | d4 |
| GF-d4-Plate1-25208-67 | 25208 | GF | Plate1 | 67 | F | 9 | 1551867 | d4 |
| WJL-d4-Plate2-25210-70 | 25210 | WJL | Plate2 | 70 | F | 6 | 8073425 | d4 |
| GF-d4-Plate1-25208-72 | 25208 | GF | Plate1 | 72 | C | 9 | 2668655 | d4 |
| GF-d4-Plate2-25210-74 | 25210 | GF | Plate2 | 74 | B | 5 | 947737 | d4 |
| WJL-d4-Plate2-25210-75 | 25210 | WJL | Plate2 | 75 | C | 6 | 4812520 | d4 |
| GF-d4-Plate2-25183-78 | 25183 | GF | Plate2 | 78 | B | 1 | 2869820 | d4 |
| WJL-d4-Plate2-25183-79 | 25183 | WJL | Plate2 | 79 | C | 2 | 4934533 | d4 |
| GF-d4-Plate1-25210-83 | 25210 | GF | Plate1 | 83 | C | 5 | 4113175 | d4 |
| WJL-d4-Plate1-25210-84 | 25210 | WJL | Plate1 | 84 | B | 6 | 4684552 | d4 |
| GF-d4-Plate2-25208-86 | 25208 | GF | Plate2 | 86 | B | 9 | 3324070 | d4 |
| GF-d4-Plate1-25183-87 | 25183 | GF | Plate1 | 87 | C | 1 | 3728767 | d4 |
| WJL-d4-Plate1-25183-88 | 25183 | WJL | Plate1 | 88 | B | 2 | 4564509 | d4 |
| WJL-d4-Plate1-25210-90 | 25210 | WJL | Plate1 | 90 | C | 6 | 3714293 | d4 |
| GF-d4-Plate1-25210-91 | 25210 | GF | Plate1 | 91 | B | 5 | 4179985 | d4 |
| GF-d4-Plate2-25208-93 | 25208 | GF | Plate2 | 93 | C | 9 | 3569201 | d4 |
| WJL-d4-Plate1-25183-94 | 25183 | WJL | Plate1 | 94 | C | 2 | 4200621 | d4 |
| GF-d4-Plate1-25183-95 | 25183 | GF | Plate1 | 95 | B | 1 | 4373035 | d4 |
| GF-d4-Plate1-25208-98 | 25208 | GF | Plate1 | 98 | B | 9 | 3652231 | d4 |
| WJL-d4-Plate2-25210-101 | 25210 | WJL | Plate2 | 101 | B | 6 | 4457721 | d4 |
| GF-d4-Plate2-25210-103 | 25210 | GF | Plate2 | 103 | C | 5 | 3903565 | d4 |
| WJL-d4-Plate2-25183-104 | 25183 | WJL | Plate2 | 104 | B | 2 | 982388 | d4 |
| GF-d4-Plate2-25183-105 | 25183 | GF | Plate2 | 105 | C | 1 | 3094592 | d4 |
| GF-d4-Plate2-25208-110 | 25208 | GF | Plate2 | 110 | A | 9 | 1967561 | d4 |
| WJL-d4-Plate1-25210-112 | 25210 | WJL | Plate1 | 112 | A | 6 | 3472086 | d4 |
| WJL-d4-Plate1-25183-116 | 25183 | WJL | Plate1 | 116 | A | 2 | 4865847 | d4 |
| GF-d4-Plate2-25210-119 | 25210 | GF | Plate2 | 119 | A | 5 | 3773438 | d4 |
| WJL-d4-Plate2-25208-120 | 25208 | WJL | Plate2 | 120 | F | 10 | 2018688 | d4 |
| WJL-d4-Plate2-25208-121 | 25208 | WJL | Plate2 | 121 | D | 10 | 2595705 | d4 |
| WJL-d4-Plate2-25208-123 | 25208 | WJL | Plate2 | 123 | E | 10 | 1841390 | d4 |
| WJL-d4-Plate2-25208-124 | 25208 | WJL | Plate2 | 124 | A | 10 | 3326544 | d4 |
| GF-d4-Plate2-25183-125 | 25183 | GF | Plate2 | 125 | A | 1 | 1822797 | d4 |
| WJL-d4-Plate2-25208-126 | 25208 | WJL | Plate2 | 126 | B | 10 | 3831425 | d4 |
| WJL-d4-Plate2-25208-127 | 25208 | WJL | Plate2 | 127 | C | 10 | 3109485 | d4 |
| WJL-d4-Plate2-25210-129 | 25210 | WJL | Plate2 | 129 | A | 6 | 1737064 | d4 |
| GF-d4-Plate1-25208-132 | 25208 | GF | Plate1 | 132 | A | 9 | 3284211 | d4 |
| WJL-d4-Plate2-25183-135 | 25183 | WJL | Plate2 | 135 | A | 2 | 4603643 | d4 |
| GF-d4-Plate1-25210-139 | 25210 | GF | Plate1 | 139 | A | 5 | 2749602 | d4 |
| GF-d4-Plate1-25183-140 | 25183 | GF | Plate1 | 140 | A | 1 | 2722703 | d4 |

## References

1. Flatt, T. (2005). The evolutionary genetics of canalization. Q Rev Biol 80, 287–316.

2. Rutherford, S., Hirate, Y., and Swalla, B.J. (2007). The Hsp90 capacitor, developmental remodeling, and evolution: the robustness of gene networks and the curious evolvability of metamorphosis. Crit Rev Biochem Mol Biol 42, 355–372.

3. Wagner, A. (2007). Robustness and Evolvability in Living Systems., Paperback Edition, (Princeton University Press).

4. Brucker, R.M., and Bordenstein, S.R. (2013). The hologenomic basis of speciation: gut bacteria cause hybrid lethality in the genus Nasonia. Science 341, 667–669.

5. Rosenberg, E., and Zilber-Rosenberg, I. (2011). Symbiosis and development: the hologenome concept. Birth Defects Res C Embryo Today 93, 56–66.

6. Douglas, A.E. (2014). Symbiosis as a general principle in eukaryotic evolution. Cold Spring Harb Perspect Biol 6.

7. Storelli, G., Strigini, M., Grenier, T., Bozonnet, L., Schwarzer, M., Daniel, C., Matos, R., and Leulier, F. (2017). Drosophila Perpetuates Nutritional Mutualism by Promoting the Fitness of Its Intestinal Symbiont Lactobacillus plantarum. Cell metabolism.

8. Storelli, G., Defaye, A., Erkosar, B., Hols, P., Royet, J., and Leulier, F. (2011). Lactobacillus plantarum promotes Drosophila systemic growth by modulating hormonal signals through TOR-dependent nutrient sensing. Cell metabolism 14, 403–414.

9. Erkosar, B., Storelli, G., Mitchell, M., Bozonnet, N., Leulier, F. (2015). Pathogen virulence impedes mutualist-mediated enhancement of host protein digestion capaciity and juvenile growth. in review.

10. Lehner, B. (2013). Genotype to phenotype: lessons from model organisms for human genetics. Nature reviews. Genetics 14, 168–178.

11. Alpern, D., Gardeux, V., Russeil, J., and Deplancke, B. (2018). Time-and cost-efficient high-throughput transcriptomics enabled by Bulk RNA Barcoding and sequencing. bioRxiv.

12. Erkosar, B., Defaye, A., Bozonnet, N., Puthier, D., Royet, J., and Leulier, F. (2014). Drosophila microbiota modulates host metabolic gene expression via IMD/NF-kappaB signaling. Plos one 9, e94729.

13. Mestek Boukhibar, L., and Barkoulas, M. (2016). The developmental genetics of biological robustness. Annals of botany 117, 699–707.

14. Posadas, D.M., and Carthew, R.W. (2014). MicroRNAs and their roles in developmental canalization. Current opinion in genetics & development 27, 1–6.

15. Tefit, M.A., Gillet, B., Joncour, P., Hughes, S., and Leulier, F. (2017). Stable association of a Drosophila-derived microbiota with its animal partner and the nutritional environment throughout a fly population’s life cycle. J Insect Physiol.

16. Rohner, N., Jarosz, D.F., Kowalko, J.E., Yoshizawa, M., Jeffery, W.R., Borowsky, R.L., Lindquist, S., and Tabin, C.J. (2013). Cryptic variation in morphological evolution: HSP90 as a capacitor for loss of eyes in cavefish. Science 342, 1372–1375.

17. Rutherford, S.L., and Lindquist, S. (1998). Hsp90 as a capacitor for morphological evolution. Nature 396, 336–342.

18. Queitsch, C., Sangster, T.A., and Lindquist, S. (2002). Hsp90 as a capacitor of phenotypic variation. Nature 417, 618–624.

19. Mirth, C.K., and Shingleton, A.W. (2012). Integrating body and organ size in Drosophila: recent advances and outstanding problems. Frontiers in endocrinology 3, 49.

20. Waddington, C.H. (1959). Canalization of development and genetic assimilation of acquired characters. Nature 183, 1654–1655.

21. Santabarbara-Ruiz, P., Lopez-Santillan, M., Martinez-Rodriguez, I., Binagui-Casas, A., Perez, L., Milan, M., Corominas, M., and Serras, F. (2015). ROS-Induced JNK and p38 Signaling Is Required for Unpaired Cytokine Activation during Drosophila Regeneration. Plos genetics 11, e1005595.

22. Jones, R.M., Luo, L., Ardita, C.S., Richardson, A.N., Kwon, Y.M., Mercante, J.W., Alam, A., Gates, C.L., Wu, H., Swanson, P.A., et al. (2013). Symbiotic lactobacilli stimulate gut epithelial proliferation via Nox-mediated generation of reactive oxygen species. EMBO J 32, 3017–3028.

23. Jones, R.M., Desai, C., Darby, T.M., Luo, L., Wolfarth, A.A., Scharer, C.D., Ardita, C.S., Reedy, A.R., Keebaugh, E.S., and Neish, A.S. (2015). Lactobacilli Modulate Epithelial Cytoprotection through the Nrf2 Pathway. Cell Rep 12, 1217–1225.

24. Broderick, N.A., Buchon, N., and Lemaitre, B. (2014). Microbiota-induced changes in drosophila melanogaster host gene expression and gut morphology. mBio 5, e01117–01114.

25. Wong, A.C., Chaston, J.M., and Douglas, A.E. (2013). The inconstant gut microbiota of Drosophila species revealed by 16S rRNA gene analysis. The ISME journal 7, 1922–1932.

26. Elgart, M., Stern, S., Salton, O., Gnainsky, Y., Heifetz, Y., and Soen, Y. (2016). Impact of gut microbiota on the fly’s germ line. Nature communications 7, 11280.

27. Bates, D., Mâchler M, Bolker B, Walker S (2015). Fitting Linear Mixed-Effects Models Using lme4. Journal of Statistical Software 67, 1–48.

28. Huang, W., Massouras, A., Inoue, Y., Peiffer, J., Ramia, M., Tarone, A.M., Turlapati, L., Zichner, T., Zhu, D., Lyman, R.F., et al. (2014). Natural variation in genome architecture among 205 Drosophila melanogaster Genetic Reference Panel lines. Genome research 24, 1193–1208.

29. Mackay, T.F., Richards, S., Stone, E.A., Barbadilla, A., Ayroles, J.F., Zhu, D., Casillas, S., Han, Y., Magwire, M.M., Cridland, J.M., et al. (2012). The Drosophila melanogaster Genetic Reference Panel. Nature 482, 173–178.

30. Dobin, A., Davis, C.A., Schlesinger, F., Drenkow, J., Zaleski, C., Jha, S., Batut, P., Chaisson, M., and Gingeras, T.R. (2013). STAR: ultrafast universal RNA-seq aligner. Bioinformatics 29, 15–21.

31. Law, C.W., Chen, Y., Shi, W., and Smyth, G.K. (2014). voom: Precision weights unlock linear model analysis tools for RNA-seq read counts. Genome biology 15, R29.

32. Ritchie, M.E., Phipson, B., Wu, D., Hu, Y., Law, C.W., Shi, W., and Smyth, G.K. (2015). limma powers differential expression analyses for RNA-sequencing and microarray studies. Nucleic acids research 43, e47.

33. Schneider, C.A., Rasband, W.S., and Eliceiri, K.W. (2012). NIH Image to ImageJ: 25 years of image analysis. Nature methods 9, 671–675.

34. Ryu, J.H., Kim, S.H., Lee, H.Y., Bai, J.Y., Nam, Y.D., Bae, J.W., Lee, D.G., Shin, S.C., Ha, E.M., and Lee, W.J. (2008). Innate immune homeostasis by the homeobox gene caudal and commensal-gut mutualism in Drosophila. Science 319, 777–782.

35. Hu, Y., Comjean, A., Perkins, L.A., Perrimon, N., and Mohr, S.E. (2015). GLAD: an Online Database of Gene List Annotation for Drosophila. J Genomics 3, 75–81.

36. Powell, J.A. (2014). GO2MSIG, an automated GO based multi-species gene set generator for gene set enrichment analysis. BMC Bioinformatics 15, 146.

37. Saunders, A., Werner, J., Andrulis, E.D., Nakayama, T., Hirose, S., Reinberg, D., and Lis, J.T. (2003). Tracking FACT and the RNA polymerase II elongation complex through chromatin in vivo. Science 301, 1094–1096.

38. Shimojima, T., Okada, M., Nakayama, T., Ueda, H., Okawa, K., Iwamatsu, A., Handa, H., and Hirose, S. (2003). Drosophila FACT contributes to Hox gene expression through physical and functional interactions with GAGA factor. Genes & development 17, 1605–1616.

